# Pathological brain states in Alzheimer’s disease

**DOI:** 10.1101/2023.08.30.555617

**Authors:** Jenna N. Adams, Sarah M. Kark, Miranda G. Chappel-Farley, Yuritza Escalante, Lea A. Stith, Paul E. Rapp, Michael A. Yassa, the Alzheimer’s Disease Neuroimaging Initiative

**Affiliations:** Department of Neurobiology and Behavior and Center for the Neurobiology of Learning and Memory, University of California, Irvine; Department of Military and Emergency Medicine, Uniformed Services University, Bethesda, Maryland

**Author notes:** Data used in preparation of this article were obtained from the Alzheimer’s Disease Neuroimaging Initiative (ADNI) database (adni.loni.usc.edu). As such, the investigators within the ADNI contributed to the design and implementation of ADNI and/or provided data but did not participate in analysis or writing of this report. A complete listing of ADNI investigators can be found at: http://adni.loni.usc.edu/wp-content/uploads/how_to_apply/ADNI_Acknowledgement_List.pdf. Corresponding authors: Jenna N. Adams, Michael A. Yassa.

## Abstract

Dynamic and rapid reconfigurations of neural activation patterns, known as brain states, support cognition. Recent analytic advances applied to functional magnetic resonance imaging now enable the quantification of brain states, which offers a substantial methodological improvement in characterizing spatiotemporal dynamics of activation over previous functional connectivity methods. Dysfunction to the persistence and temporal transitions between discrete brain states may be proximal factors reflecting neurophysiological disruptions in Alzheimer’s disease, although this has not yet been established. Here, we identified six distinct brain states, representing spatiotemporal trajectories of coactivation at single time points, in older adults across the Alzheimer’s disease continuum. Critically, we identified a pathological brain state that reflects coactivation within limbic regions. Higher persistence within and transitions to this limbic state, at the expense of other brain states, is associated with an increased likelihood of a clinically impaired diagnosis, worse cognitive performance, greater Alzheimer’s pathology, and neurodegeneration. Together, our results provide compelling evidence that neural activity settling into a pathological limbic state reflects the progression to Alzheimer’s disease. As brain states have recently been shown to be modifiable targets, this work may inform the development of novel neuromodulation techniques to reduce limbic state persistence. This application would be an innovative clinical approach to rescue cognitive decline in the early stages of Alzheimer’s disease.

## INTRODUCTION

The brain is a dynamical system – rapidly responding to external cues and internal goals. Coordinated activation within cortical networks is a highly dynamic process, with specialized networks coming online and interacting in a fluid manner that is related to cognitive and task demands^2,8–11^. These transient whole-brain patterns of coactivation, also known as *brain states* (for a review, see Greene et al., 2023^1^), can now be evaluated non-invasively in humans with functional magnetic resonance imaging in combination with sophisticated modeling techniques^2,8,12–15^. This temporal modeling of regional activation space enables vastly richer information reflecting dynamics of networks compared to what can be measured with traditional measures of static functional connectivity or sliding time window approaches, both of which reduce the time dimension and simplify complex interactions among brain regions as pairwise correlations^3,4^.

While the dynamic nature of the brain provides flexibility and the ability for differential processing, it also may confer an inherent vulnerability if these dynamics become dysfunctional. In many neuropsychiatric conditions (e.g. major depressive disorder and schizophrenia^6,16–20^) and developmental disorders (e.g. autism spectrum disorder^21^), the brain has been shown to persist (or *dwell*) in certain patterns of activation, settling into potentially pathological low-energy states. Further, the temporal trajectory of how the brain traverses these states can also become dysfunctional, with changes to the overall frequency of transitions or probability of transitioning between certain states. These differences in persistence and transition probabilities are associated with maladaptive behavioral responses, and interventional techniques to manipulate brain states have been found to improve function^22^.

Time-varying dynamics of brain states have not yet been characterized in neurodegenerative diseases such as Alzheimer’s disease. Alzheimer’s disease and related dementias are estimated to affect over 55 million people worldwide, and this number is projected to rapidly increase in the coming years as the aging population increases^23^. Alzheimer’s disease is characterized by a pathological cascade involving accumulation of aggregated proteins^24^, neurophysiological disruptions^25,26^, and neurodegeneration^27^, which together express as memory and cognitive impairment^28^ (for a review, see Knopman et al., 2021^29^). It is well known that aging and Alzheimer’s disease is associated with neurophysiological changes such as neuronal hyperexcitability^25^, impaired excitatory-inhibitory balance^30^, and abnormal static functional connectivity between regions^31^ and large-scale cortical networks^32,33^.

In contrast to the extensive characterization of functional networks in aging and Alzheimer’s disease using static metrics of functional networks^31–33^, and to a lesser extent, dynamic functional connectivity approaches with sliding time windows^34–39^, time-varying dynamics of brain states, specifically reflecting coactivation patterns at single time points, have not yet been investigated. Insight into the rapid spatiotemporal dynamics of brain states would provide a more precise understanding of the mechanism underlying *how* network dysfunction may emerge. For example, using brain state analyses, we can probe whether there are specific coactivation patterns, or pathological “states”, in which the aging brain transitions to and dwells within that are related to neuropathology and cognitive dysfunction. As brain states have recently been shown to be modifiable^5,6^, these pathological states may be ideal targets for intervention. Thus, the characterization of brain state dynamics in older adults is critical as it may offer a mechanistic, modifiable link between the transition from healthy aging to the onset of Alzheimer’s disease.

In the current study, we characterized brain state dynamics in a sample of older adults ranging from cognitively unimpaired to patients with mild cognitive impairment and dementia. We leverage resting state fMRI with high spatial and temporal resolution combined with a rich dataset of clinical, cognitive, and neuropathological measures of Alzheimer’s disease (Alzheimer’s Disease Neuroimaging Initiative, ADNI3^40^) to interrogate relationships between brain states and dysfunction related to the onset of Alzheimer’s disease. We demonstrate that increased persistence and transitions to a brain state reflecting limbic coactivation is associated with worse clinical outcomes, greater Alzheimer’s disease pathology and neurodegeneration, and worse cognitive performance, suggesting time varying dynamics of brain states are a key contributor to Alzheimer’s disease pathophysiology.

### Six distinct brain states represent spatiotemporal coactivation patterns in older adults

To investigate how brain states may relate to the progression from healthy aging to Alzheimer’s disease, we analyzed 334 high resolution resting state fMRI scans from 201 participants in the Alzheimer’s Disease Neuroimaging Initiative (ADNI3^40^), spanning from cognitively unimpaired (n=188 scans, 56.3%), mild cognitive impairment (n=115 scans, 34.4%), and dementia (n=31 scans, 9.3%) (age range 49-97, mean 73.9 years old, 50.9% female; for further demographic information, see **Extended Data Table 1**; **Methods**). To derive dynamic brain states from resting state fMRI data, we performed k-means clustering on the mean BOLD time series from 218 cortical regions of interest^2^ (see **Methods**). A clustering solution corresponding to six brain states was selected using data-driven approaches (**Extended Data Figure 1**). These brain states, depicted in **Figure 1a**, represent distinct patterns of high and low amplitude coactivation at single time points that consistently co-occur across participants and the course of the scan. Critically, brain state centroids were not influenced by potential confounding factors such as motion or scan ramp-up effects (see **Methods**) and demonstrated high split-half reliability and robustness compared to several null data models (**Extended Data Figures 2 and 3**). Each spatial pattern of coactivation (**Figure 1a**) was compared to canonical resting state networks^41^ using cosine similarity to facilitate interpretation of each state (**Figure 1b**).

**Figure 1.**
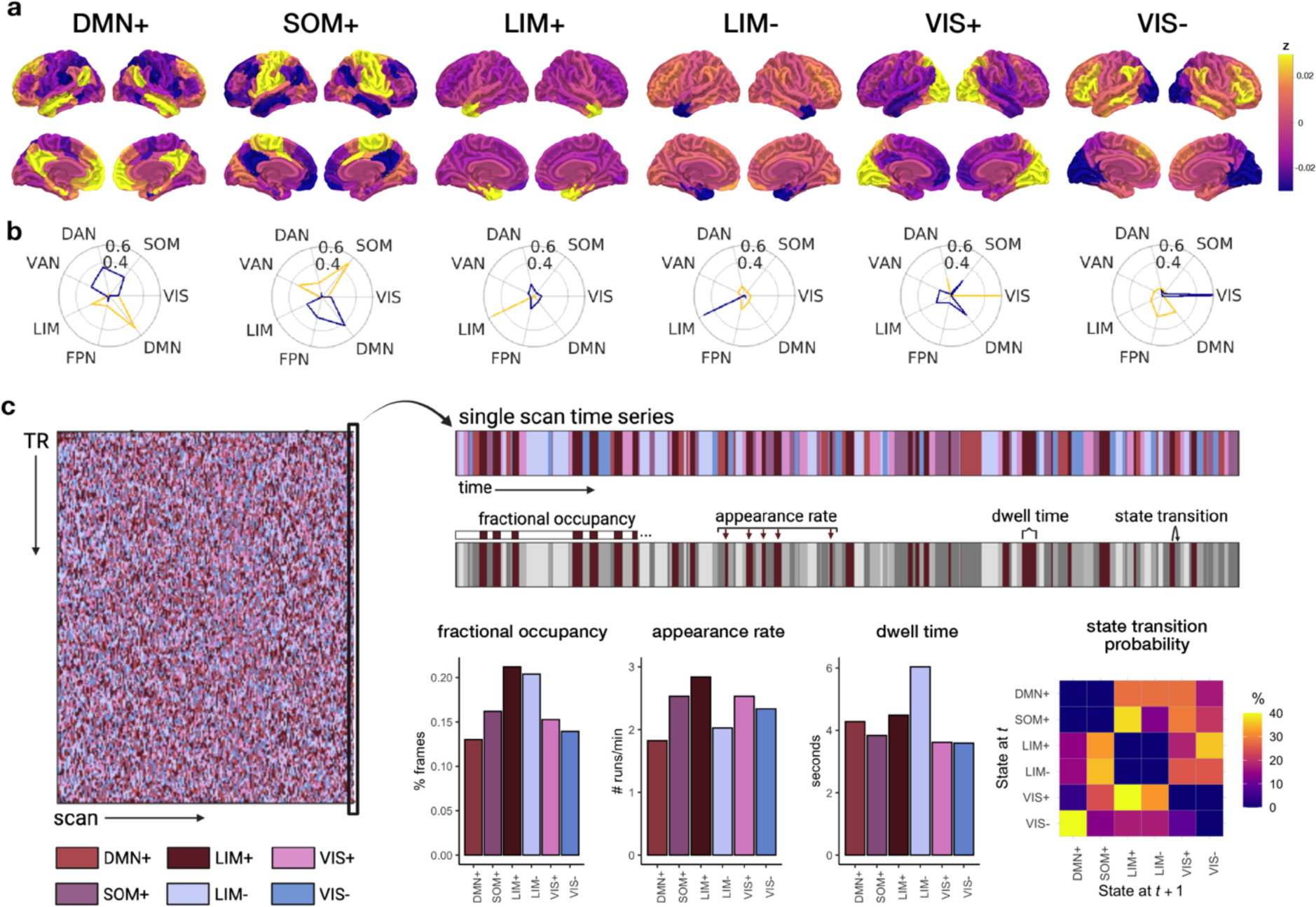
Visualization of brain states and derivation of brain state persistence features and transition probabilities. (**a**) Six distinct brain states reflecting spatiotemporal patterns of whole brain coactivation were identified with a k-means clustering approach. The spatial pattern of coactivation in each brain state was compared to canonical resting state networks using cosine similarity, depicted in the polar plots in (**b**) The outer circle boundary represents cosine similarity of 0.6, while the inner circle represents a cosine similarity value of 0.4. The resting state network with the more similar high amplitude (yellow, +) or low amplitude (blue, −) coactivation pattern was used to assign each brain state a name in (**a**), however, brain states tend to reflect coactivation across resting state networks. (**c**) In the k-means clustering approach, each time point (TR, y-axis) of each scan (x-axis) was assigned to a state. For a single scan’s time series, the progression of states can be represented over time (top right). We characterized three persistence features for each state, demonstrated for the LIM+ state in the example scan’s time series: fractional occupancy, or the overall percentage of time spent in that state; appearance rate, or how often a state occurred per minute; and dwell time, or how long on average a state persisted in seconds. We also characterized state transition probability from each state at time *t* to the state at time point *t + 1*. DMN+, default mode network high amplitude state; SOM+, somatomotor network high amplitude state; LIM+, limbic network high amplitude state; LIM-, limbic network low amplitude state; VIS+, visual network high amplitude state; VIS-, visual network low amplitude state; DAN, dorsal attention network, DMN, default mode network, FPN, frontoparietal network, LIM, limbic network, SOM, somatomotor network; VAN, ventral attention network; VIS, visual network.

Six brain states were identified that corresponded to patterns of: (1) high amplitude default mode network coactivation, with secondary low amplitude dorsal attention, ventral attention, and somatomotor network coactivation (DMN+ state); (2) high amplitude coactivation in somatomotor regions, with secondary low amplitude coactivation in default mode network regions (SOM+ state); (3) high amplitude limbic coactivation, with minimal and non-specific low amplitude coactivation (LIM+ state); (4) low amplitude limbic coactivation, with minimal and non-specific high amplitude coactivation (LIM-state); (5) high amplitude visual network coactivation, with secondary low amplitude coactivation of default mode and limbic networks (VIS+ state); and (6) low amplitude visual network coactivation, with secondary high amplitude coactivation of default mode and frontoparietal networks (VIS-state). Interestingly, each primary brain state component had a high amplitude and low amplitude version, with DMN+/SOM+, LIM+/LIM-, and VIS+/VIS-patterns resembling inverse spatial patterns.

As part of the brain state clustering solution, each time point (TR) across the scan was assigned to one of the six brain states (**Figure 1c**). For each scan, we quantified three persistence features of brain states (**Figure 1c**; see **Methods**): *fractional occupancy*, or the percentage of the scan occupied by the state; *appearance rate*, or how often the state occurred on average per minute; and *dwell time*, or how long on average the state persisted once it was entered. Group average values and correlations of persistence features between states are depicted in **Extended Data Figure 4**. We also modeled the transition probability, or the likelihood of transitioning from one state to another state (**Figure 1c**, see **Methods**). Using this dynamic information from our identified brain states, we next aimed to determine how these persistence features and transition probabilities reflected clinical outcomes, cognitive performance, and Alzheimer’s disease pathophysiology.

### Brain state persistence features predict clinical and cognitive outcomes

To test whether persistence features of brain states provided sensitive and meaningful information about clinical and cognitive outcomes, we entered all 18 persistence features (fractional occupancy, appearance rate, and dwell time for each state) into a second level of k-means clustering to determine brain state *profiles*, or common patterns of brain state persistence features across subjects. Four distinct brain state profiles emerged, with unique combinations of high, medium, or low levels of each feature (**Figure 2a**; see **Extended Data Figure 5** for selection of *k*=4 profiles). For example, Profile 1 demonstrated high DMN+/SOM+/VIS state and low LIM state persistence features, while in contrast, Profile 4 demonstrated high LIM state and low DMN+/SOM+/VIS state persistence features. These profiles reliably emerged across different clustering methods and significantly differed from shuffled data (**Extended Data Figure 5**).

**Figure 2.**
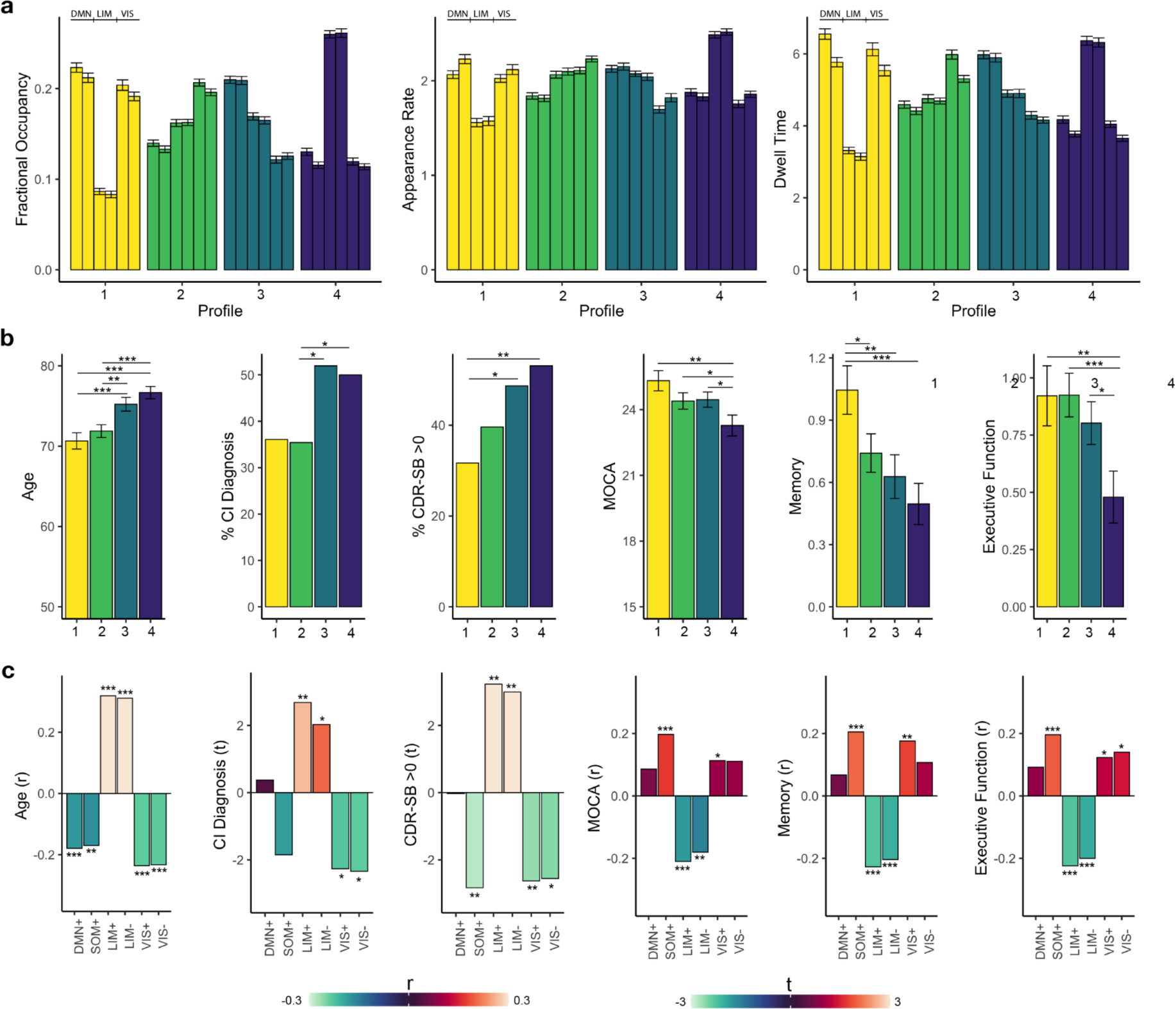
Associations between state persistence features and clinical and cognitive outcomes. (**a**) Using all 18 persistence features representing the six brain states, we derived four brain state profiles, or common patterns of brain state persistence features across scans. Within each profile, values of each state are represented as bars in the following order: DMN+, SOM+, LIM+, LIM-, VIS+, VIS-. (**b**) Profile 4 (purple) and Profile 3 (blue) had older age, worse clinical outcomes, and worse cognitive performance compared to Profile 1 (yellow) and Profile 2 (green). (**c**) Direct correlations between fractional occupancy of each state with clinical and cognitive outcomes revealed that higher LIM state fractional occupancy was the most strongly associated with worse outcomes, and may be driving results of the profile analysis in (**b**). ***p<0.001 **p<0.01 *p<0.05

We next tested whether clinical and cognitive outcomes differed across brain state profiles with distinct feature loadings (**Figure 2b**). Profiles significantly differed by age (F(3) = 10.9, p<0.001), likelihood of a clinically impaired diagnosis (cognitively unimpaired vs. MCI or dementia; X^2^(3) = 7.86, p=0.049), CDR-SB score indicating clinical dysfunction (CDR-SB >0 vs. 0; X^2^(3) = 8.36, p=0.04), and MoCA score (F(3) = 3.80, p=0.01). Further, profiles significantly differed across neuropsychological test performance in composite domains of memory (F(3) = 4.61, p=0.004) and executive function (F(3) = 4.14, p=0.007). Planned follow-up pairwise comparisons revealed that these differences were driven by impairment in Profiles 3 and 4, which demonstrated older age, more clinical impairment, and worse cognitive performance compared to Profiles 1 and 2 (**Figure 2b**; see **Extended Data Table 2** for pairwise comparison statistics). Profile 1, characterized by low LIM persistence and high persistence of other states, had the healthiest outcomes compared to other profiles. In contrast, Profile 4, characterized by high LIM persistence and low persistence of other states, consistently had the worst outcomes over the tested measures, and even demonstrated a significantly lower MoCA and memory composite score compared to Profile 3 (**Figure 2b; Extended Data Table 1**). These results indicated that Profile 4 may reflect a dysfunctional combination of brain states that contributes to poor clinical and cognitive outcomes.

To further determine which brain states were most strongly contributing to poor outcomes, we next tested direct relationships between the fractional occupancy of each state and outcomes (**Figure 2c**). We found that increased fractional occupancy of the LIM+/LIM-states, and to a lesser extent, decreased fractional occupancy of the remaining states, was associated with worse outcomes (**Figure 2c**), confirming that high persistence of LIM states may be driving the differences between the profiles. Consistent with this finding, Profile 1, the healthiest profile in relation to behavioral and diagnostic outcomes, had the lowest LIM state persistence, while Profile 4, the most impaired profile, had the highest LIM state persistence.

### Increased transition probability to limbic states reflects dysfunction

To further characterize temporal dynamics of brain states, we modeled the probability of transitions between brain states from the state at time *t* to *t+1* (see **Methods**). There was a higher probability of remaining in the same state compared to switching between states, consistent with previous work^5^. To better understand trajectories between distinct states, we modeled transition probabilities after removing the effects of autocorrelation to determine transition trajectories without influence of persistence features^2^ (**Figure 3a-b**; see **Methods**). At the group level, transitioning from the SOM+ state to the LIM+ state (29.4%), and from the DMN+ state to the LIM-state (30.0%), had the highest probability, while transitioning from a high to low amplitude version of the same state was the least likely (0.03%-4.5% range).

**Figure 3.**
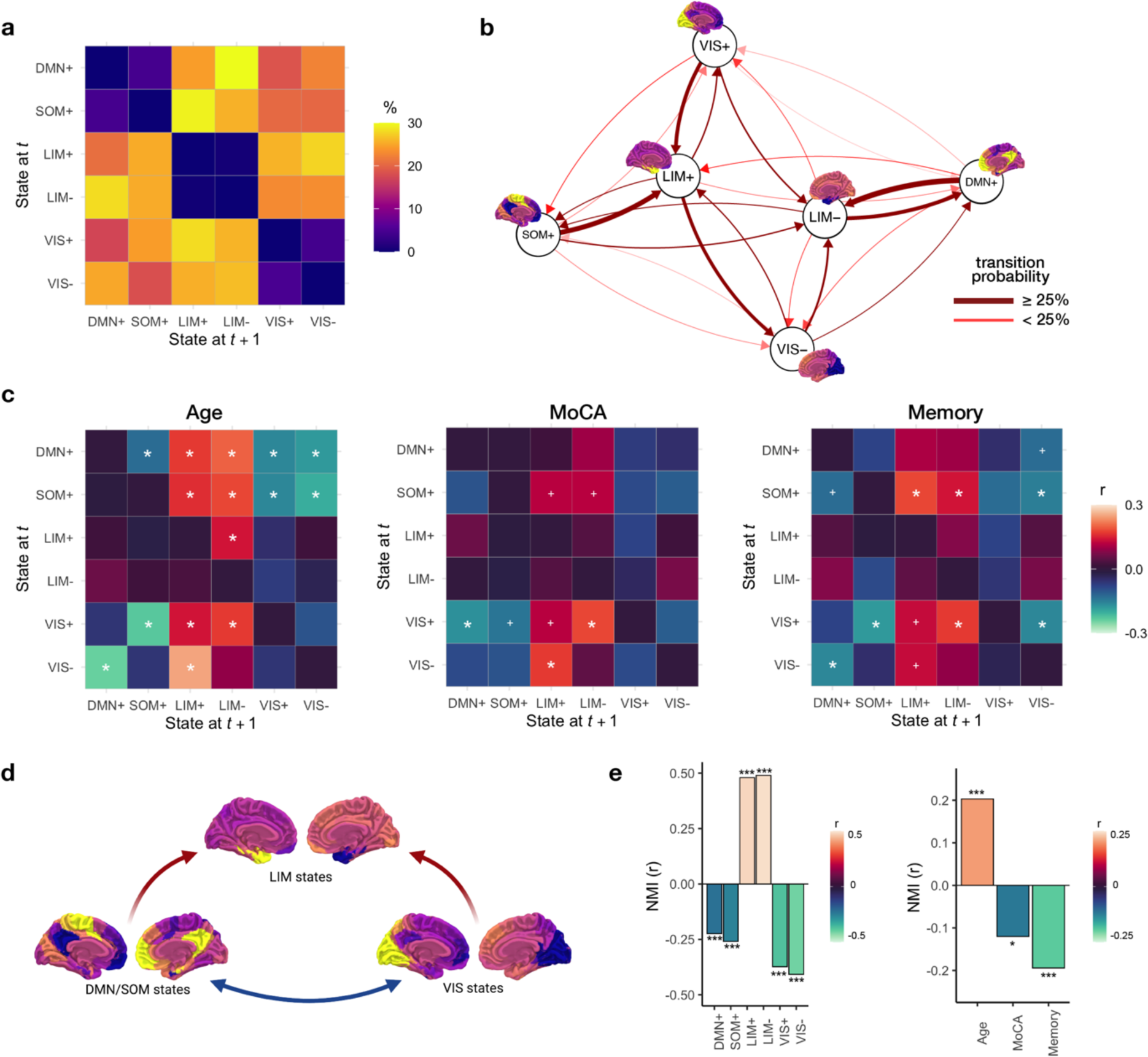
State transition probability and associations with clinical and cognitive outcomes. (**a**) Group-average transition matrix showing the probability (percentage) of transitioning from a state at time *t* to a state at time *t+1*. (**b**) Transition probability results from (**a**) represented as a flow diagram. The highest probability transitions (>25%) are shown as thicker dark red lines, moderately probable transitions (<25%) are shown as thinner red lines. Transitions with <5% likelihood (representing transitions from high to low amplitude versions of the same state, or vice versa) are not depicted. (**c**) Associations between age, clinical impairment (MoCA score) and memory performance with state transitions. Correlation coefficients are represented as positive (reds) or negative (blues) between each transition probability and outcome measure. Significance was corrected for multiple comparisons using FDR correction over 30 possible transitions. White asterisks represent pFDR <0.05, and white crosses represent punc <0.05. (**d**) Schematic diagram depicting interpretation of results shown in (**c**). Higher probability of transitions to the LIM states from DMN+/SOM+ or VIS+/VIS-states (red arrows), and lower probability of transitions between DMN+/SOM+ and VIS+/VIS-states (blue arrows) are associated with worse clinical and cognitive outcomes. (**e**) Normalized mutual information (NMI), representing the information held about future states from the current state, is related to greater LIM state prevalence and less DMN+/SOM+ or VIS state prevalence. Greater NMI is also associated with older age, lower MoCA score, and worse memory performance.

We next tested whether state transition probabilities were associated with clinical and cognitive outcomes. An increased probability of transitioning into LIM+/LIM-states from DMN+/SOM+ or VIS+/VIS-states, and a decreased probability of transitioning between DMN+/SOM+ and VIS+/VIS-states, was strongly associated with increased age, greater clinical impairment, and worse memory performance (psFDR<0.05; **Figure 3c**; see **Extended Data Figure 6** for additional outcome variables). Transition probability out of the LIM+/LIM-states to other states were not strongly associated with outcomes. Together, these state transition patterns suggest that an increased probability of entering into a LIM+/LIM-state at the cost of transitioning to a DMN-/SOM+ or VIS+/VIS-state is associated with poor clinical and cognitive outcomes (**Figure 3d**), consistent with the brain settling into a dysfunctional low energy LIM state.

Finally, we tested the degree to which the current brain state held information about transitioning to the next brain state with a measure of normalized mutual information (NMI) between lagged state time series (see **Methods**). Greater NMI was associated with greater fractional occupancy of the LIM states, and lower fractional occupancy of the DMN+/SOM+ and VIS states (ps<0.001; **Figure 3e**). Further, greater NMI was associated with older age (r = 0.20, p<0.001), lower MoCA score (r = −0.12, p = 0.04), and lower memory performance (r = −0.19, p<0.001; **Figure 3e**). These results are consistent in suggesting that increased persistence of LIM states and poor outcomes are associated with a more predictable brain state trajectory that is characterized by greater transitions to LIM states.

### Limbic state persistence features are related to Alzheimer’s neuropathology

The development of pathology and neurodegeneration are proximal factors underlying clinical and cognitive outcomes in Alzheimer’s disease^29^. We next tested whether brain state persistence features were related to these pathological changes. To assess this, we conducted a sparse canonical correlation analysis (see **Methods**) to test whether brain state persistence features were associated with regionally-specific spatial patterns of Alzheimer’s pathology and neurodegeneration. Increased persistence of the LIM+ state and decreased persistence of the SOM+ state (**Figure 4a**) were associated with higher tau pathology in medial and inferior temporal regions (r = 0.23, p<0.001; **Figure 4b**), and higher Aβ pathology in inferolateral and posterior occipital regions (r = 0.18, p = 0.006; **Figure 4b**), reflecting patterns of known regional vulnerability to pathology. A combination of increased LIM+ state persistence and decreased VIS+/VIS-state persistence (**Figure 4a**) was associated with decreased volume in regions overlapping with pathology, particularly in medial and lateral temporal lobe, and medial and lateral parietal lobes (r = 0.36, p<0.001; **Figure 4b**).

**Figure 4.**
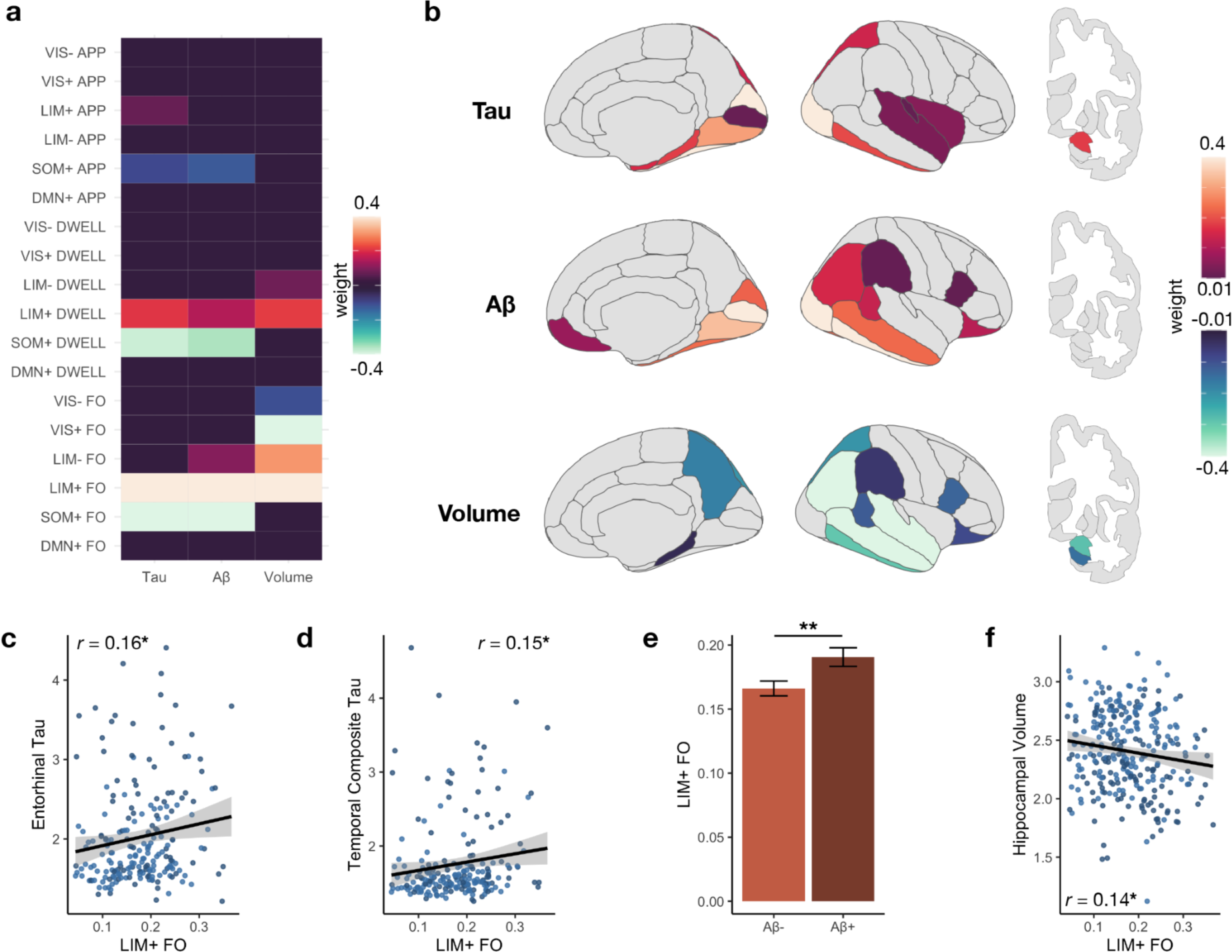
Associations between brain state persistence features and Alzheimer’s disease neuropathology. To determine spatial patterns of tau, Aβ, and volume that reflected brain state persistence features, we conducted sparse canonical correlation analyses. Weight loadings for brain state persistence features are shown in (**a**), with their associated spatial pattern rendered on brain images in (**b**). Increased LIM+ state persistence (hot colors) and decreased SOM+/VIS+ state persistence (cool colors) were significantly associated with greater tau and Aβ deposition (hot colors) and decreased regional volume (cool colors) in regionally specific patterns mirroring known vulnerability. A priori investigations of (**c**) entorhinal tau, (**d**) temporal composite tau, (**e**) Aβ positivity status, and (**f**) hippocampal volume revealed strong associations with LIM+ fractional occupancy. Scatterplots (**c, d, f**) show cognitively unimpaired in lighter blue circles, and cognitively impaired in darker blue circles. FO, fractional occupancy; DWELL, dwell time; APP, appearance rate; *p<0.05, **p<0.01

Increases in LIM+ state fractional occupancy showed the strongest and most consistent associations with tau, Aβ pathology, and volume using the data-driven approach, while decreases in SOM+ and VIS state persistence contributed to pathology and neurodegeneration, respectively. To further probe our data-driven results, we next conducted targeted analyses with *a priori* selected outcome measures of pathology. We focused on tau pathology within entorhinal cortex, the first cortical region to accumulate tau pathology in aging and preclinical AD^24^, and within a temporal composite region, a set of regions previously shown to be sensitive to AD-related tau progression^42^. Increased LIM+ state fractional occupancy was associated with higher tau in entorhinal cortex (r = 0.16, p = 0.02; **Figure 4c**), and in the temporal composite (r = 0.15, p = 0.035; **Figure 4d**), Within cognitively unimpaired older adults, higher LIM+ fractional occupancy was more strongly associated with entorhinal tau (r = 0.21, p = 0.026), while lower SOM+ fractional occupancy was associated with temporal composite tau (r = −0.24, p = 0.009). These relationships did not persist in older adults with cognitive impairment (MCI/Dementia; entorhinal & LIM+ fractional occupancy: r = 0.06, p = 0.56; temporal composite tau & SOM+ fractional occupancy: r = −0.13, p = 0.24), suggesting these states may more strongly contribute to or emerge from the early stages of tau accumulation. To characterize Aβ burden, we compared older adults classified as Aβ-versus Aβ+ using a validated threshold and set of composite regions^43,44^. Older adults who were Aβ+ had higher fractional occupancy of the LIM+ state (t(226) = −2.67, p = 0.008; **Figure 4e**). Again, this relationship remained significant within the cognitively unimpaired older adults (t(133) = −2.09, p = 0.039), but not within cognitively impaired older adults (t(91) = −0.56, p = 0.58). Together, these results suggest that increased LIM+ fractional occupancy is highly sensitive to pathology in the prodromal phase of Alzheimer’s disease.

We next investigated direct associations with hippocampal volume as a sensitive marker for Alzheimer’s-related neurodegeneration^45^. Increased LIM+ fractional occupancy was associated with decreased hippocampal volume (r = −0.14, p = 0.02; **Figure 4f**). In contrast to the pathology results, relationships between LIM+ fractional occupancy and hippocampal volume were strong across the whole sample, though primarily driven by the cognitively impaired participants (r = −0.16, p = 0.08) rather than unimpaired (r = −0.03, p = 0.68) participants, supporting models of AD in which neurodegeneration follows pathology in later stages of disease. Further, across the whole sample, hippocampal volume was more strongly associated with LIM+ dwell time (r = −0.18, p = 0.003) rather than appearance rate (r = −0.01, p = 0.84), and with VIS+ fractional occupancy (r = 0.21, p<0.001) rather than SOM+ fractional occupancy (r = 0.02, p = 0.72).

Associations between transition probabilities and pathology (entorhinal tau and Aβ status) did not survive multiple comparison corrections (see **Methods**), indicating that patterns of transitions may not have as strong of a relationship with pathology compared to overall persistence of each state (**Extended Data Figure 6b**). However, lower hippocampal volume was significantly associated with more transitions from the LIM+ to LIM-state (r = −0.18, pFDR = 0.002), and less transitions from the VIS- to DMN+ state (r = 0.19, pFDR = 0.002; **Extended Data Figure 6b**).

### Limbic state persistence is a strong contributor to memory performance

To further probe how brain state features may link emerging neuropathology to expression of cognitive deficits, we constructed models combining LIM+ fractional occupancy with age and neuropathology. In linear regression models predicting to performance on the MoCA (R^2^ = 0.38, p<0.001; **Extended Data Table 3**) and the memory composite (R^2^ = 0.34, p<0.001; **Extended Data Table 3**), LIM+ fractional occupancy was a significant predictor (MoCA: p = 0.01, memory: p=0.009), even when common factors such as entorhinal tau, hippocampal volume, and age that are known to be strongly related to cognition were also included in the model. Similarly, in logistic regression models predicting to clinical impairment (CDR-SB>0; R^2^ = 0.33, p<0.001; **Extended Data Table 3**) or diagnosis (CU vs MCI/Dementia; R^2^ = 0.17, p<0.001; **Extended Data Table 3**), LIM+ fractional occupancy was a significant predictor of clinical impairment (p = 0.004) and a trending predictor of diagnosis (p = 0.07) when included in the same model with entorhinal tau, hippocampal volume, and age.

We then constructed mediation models (see **Methods**) to test potential directional relationships between these factors. We found that LIM+ fractional occupancy partially mediated the relationship between age and memory performance (indirect effect: β=-0.007, p=0.006, CI [− 0.01, −0.002], 26.7% of total effect; direct effect: β = −0.02, p=0.005, CI [-0.03, −0.01]; total effect: β = −0.03, p<0.001, CI [-0.04, −0.01]). Further, the relationship between LIM+ fractional occupancy and memory was partially mediated by entorhinal tau (indirect effect: 32.2% of total effect, β = −1.09 p=0.03, CI [-2.10, −0.09]; direct effect: β = −2.30, p = 0.002, CI [-3.78, −0.82]; total effect: β =-3.39, p<0.001, CI [-5.15, −1.63]). These results further support the link between persistence of the LIM+ state and worse memory performance in the context of aging and development of tau pathology.

### Stability and clinical trajectories of brain state persistence features

While brain state persistence features have been shown to be stable within individuals and highly heritable^14^, these data suggest that particular brain state profiles are sensitive to pathological and cognitive change associated with Alzheimer’s disease onset and progression. To comprehensively evaluate their sensitivity to clinically-meaningful change, we tested the stability of remaining assigned to the same brain state persistence profile (i.e. **Figure 2a**) over time in participants with at least two scans, comparing profile assignment at time *t* compared to profile assignment at time *t + 1* (**Extended Data Figure 7)**. The overall distribution of profile transition probabilities significantly differed from chance (X^2^ = 35.30, p<0.001), suggesting an inherent pattern of how participants stay within or transition between profiles. There was an above-chance probability of staying within the original profile assignment (ps<0.05; **Extended Data Figure 7**). The probability of staying in an unhealthy profile (Profiles 3 and 4; 61.8%, *n*=55) was greater than staying in a healthy profile (Profiles 1 and 2; 38.2%, *n*=34; Cohen’s *h* = 2.23, p = 0.02), which is consistent with the decreased likelihood of clinical or cognitive improvement in the course of aging and Alzheimer’s disease.

Next, to further understand progression across brain state profiles, we tested whether certain profile transitions were more likely than chance. From Profile 1, the healthiest profile characterized by low LIM persistence, the probability of transitioning to any of the other three profiles was equivalent (17.7%) and not greater than chance (ps>0.71), indicating that if one was to leave Profile 1, there was no preferential next profile for transition. In contrast, from Profile 4, the unhealthiest profile characterized by high LIM persistence, the probability of a transition to Profile 1 (the healthiest profile), was significantly below chance (5.7%; Cohen’s *h* = −2.63, p = 0.007), again supporting the low likelihood of improvement. Collapsing across healthy (Profiles 1 and 2) and unhealthy (Profiles 3 and 4) profiles, the probability of transitioning from a healthy to an unhealthy profile (61.4%, n=27) was higher than transitioning from an unhealthy to healthy profile (38.6%, n=17), though this comparison did not reach statistical significance (Cohen’s h = 1.51, p = 0.09) in part due to limited sample size with multiple time points.

### Correspondence with traditional measures of static functional connectivity

Traditional investigations of static functional connectivity changes in aging and Alzheimer’s disease provide complementary information to the study of brain states^33^. To help interpret our findings in the context of previous static functional connectivity analyses, we additionally conducted ROI-to-ROI functional connectivity analyses (see **Methods**). We calculated within-network connectivity, between-network connectivity, and system segregation of the seven canonical resting state cortical networks^41^ (RSNs). We then correlated fractional occupancy of each brain state with these metrics representing the RSNs (**Extended Data Figure 8**). Overall, higher LIM state fractional occupancy was associated with decreased between-network connectivity and increased system segregation of the canonical limbic RSN (pFDRs <0.05), while simultaneously associated with decreased within-network FC and decreased segregation of the other RSNs (pFDRs <0.05). This is consistent with our interpretation of the LIM+ state persistence metrics, reflecting a LIM+ state that is more segregated from the fluctuations of other networks and may act as a low energy state. In contrast, DMN+/SOM+ state fractional occupancy was overall associated with increased within-network connectivity, increased between-network connectivity, and decreased system segregation of the RSNs, while VIS+/VIS-state fractional occupancy was associated with increased within-network connectivity, decreased between-network connectivity, and increased system segregation across RSNs (psFDR <0.05; **Extended Data Figure 8**).

## DISCUSSION

Brain states, reflecting rapid spatiotemporal dynamics of cortical coactivation patterns, reflect the pathophysiology of Alzheimer’s disease and clinical outcomes. Our study has identified a pathological limbic brain state in older adults, predominantly characterized by coactivation within limbic regions at rest. We demonstrate that increased persistence and transitions to this limbic brain state, at the cost of persisting in and transitioning to default mode network and visual states, is associated with clinical impairment, Alzheimer’s neuropathology, and memory and executive function deficits. These findings provide evidence for a compelling mechanism explaining network dysfunction and neurophysiological changes in the progression towards Alzheimer’s disease, and highlights the potential for prevention of dysfunctional limbic states as an innovative treatment option.

Our results suggest that the limbic brain state may reflect a pathological attractor state, in which the brain settles into and has trouble escaping, which leads to poor outcomes. We demonstrate that higher persistence, and particularly dwell time, is associated with poor outcomes. Critically, this is driven by an altered spatiotemporal trajectory that results in increased transitions into the limbic state, as opposed to out of the limbic state. This increased limbic persistence, driven by increased transitions into the limbic state, has proximal consequences. Limbic regions, and particularly the medial and anterior temporal lobe, are critical for memory encoding, consolidation, and recollection^46^, and are particularly susceptible to the development of pathology^24^. While it may seem counterintuitive that *more* time spent in limbic states is associated with worse memory, our results are consistent with accounts of medial temporal lobe hyperactivation being maladaptive for memory performance^47–50^. Further, our findings that greater persistence of limbic states are related to pathology in a spatial pattern reflecting known vulnerability^24^ is consistent with both animal and human imaging studies demonstrating that hyperactivity patterns may have a causal role in the accumulation of pathological proteins^51–53^.

Our results provide a mechanistic framework in which existing accounts of functional network changes can be interpreted. Prior investigations of age and Alzheimer’s disease network changes based on static^31–33^ and dynamic^34–39^ functional connectivity have proposed compelling models characterized by failures, strength changes, and disconnections between networks. While the previous studies were not able to leverage the moment-to-moment reconfiguration of unconstrained brain states^1^ due to methodological limitations^3,4^, they provide complementary information to our highly dynamic coactivation approach. For example, accounts that functional disconnection between medial temporal lobe and posterior midline regions drive medial temporal lobe hyperexcitability^31^ is consistent with our finding that increases in LIM state persistence features are related to increased segregation of the canonical limbic resting state network, supporting its disconnected and dysfunctional nature. Further, our results support an imbalance between networks, demonstrating a shift of processing from default mode and visual states to a predominantly limbic state, consistent with accounts of cascading failures to large-scale networks^32^. The temporal persistence and spatiotemporal trajectories of coactivation patterns revealed in our study will provide a new foundation for future activation and functional connectivity work in aging and Alzheimer’s disease.

The current study derived brain states from data acquired while participants were at “rest”, which, counterintuitively to its name, is a highly dynamic process characterized by spontaneous mental states such as mind wandering, future planning, retrospection, and visual imagery^54^, mapping onto default mode network, limbic, and visual states we observed. It is important to note that the six states identified here are not a discrete measure of all possible states the brain can visit during rest, and we focused on states that commonly co-occurred between participants. Using resting state, as opposed to task-based fMRI, as a method to capture dysfunctional brain state dynamics has tangible advantages, as resting state acquisition can be harmonized across research or clinical sites. Critically, resting state is feasible to acquire in patient populations in which performing a cognitive task in the scanner may range from difficult to impossible.

Future investigations in older adults should additionally test how brain state dynamics shift while performing different tasks, as well as reconfiguration of states between rest and task^2,55^. During active tasks, different brain states (such as frontoparietal network, dorsal/ventral attention networks, salience networks, etc.) related to those particular task goals may appear, and “dysfunctional” states and transitions may shift to networks opposed to task goals^56,57^. Critically, because brain states are modeled moment-to-moment, brain state analyses are an optimal method to reflect the human conscious experience, where real time changes in behavior could be precisely mapped to real time changes in neural dynamics. Further, the study of brain states is not limited to functional MRI, but can be applied to any rich dataset with dynamical spatial and temporal sampling of the brain, such as electroencephalography (EEG)^1^. Future work combining the precise spatial information gained from functional MRI with the fast-sampling rate of EEG may reveal more insight into time-varying dynamics and state transitions, especially when mapped to contemporaneous behavioral tasks.

The vast majority of clinical interventions for Alzheimer’s disease have focused on preventing or reducing proteinopathies with immunotherapies, with little success to date of restoring cognitive performance while simultaneously risking harmful cerebrovascular side effects^58,59^. In contrast, targeting the dynamics of large-scale cortical networks may have more proximal relationships to restoring cognitive function, and be used as a complementary approach to current anti-pathology treatments. Our finding of a dysfunctional limbic state, reflecting higher persistence and transitions, strongly suggests that a modifiable treatment strategy to reduce occurrence of this limbic state may ameliorate some aspects of cognitive decline. Brain states can be dynamically modulated with strategies to input energy into specific nodes in the system, with *in silico* models changing between states such as sleep and wakefulness^5^ and pharmacological manipulations demonstrating changes in the amount of energy needed to switch states^6^. In parallel to closed-loop stimulation studies, in which neurofeedback dynamically regulates disorders such as temporal lobe epilepsy^60^, brain states could be conceptualized to be modified in a similar fashion, with *brain state dependent brain stimulation*^61^ being applied when dysfunctional states are identified. Further, non-invasive neuromodulation techniques, such as transcranial magnetic stimulation^62,63^, vagus nerve stimulation^64^, closed-loop neurofeedback using neuroimaging^65^, and gamma-band entrainment^66^ (but see also Soula et al., 2023^67^), may offer other avenues to directly manipulate brain states. While still speculative and highly experimental^7^, future studies should take concrete steps to determine if manipulating brain states would in fact be a viable strategy.

In summary, we provide the first comprehensive investigation of brain states, representing spatiotemporal dynamics of coactivation at single timepoints, in the context of the progression from aging to Alzheimer’s disease. We provide a mechanistic account of how dysfunctional dynamics, namely increased persistence and transitions to a pathological state reflecting limbic coactivation, may be a proximal factor to the expression of clinical impairment, Alzheimer’s neuropathology, and cognitive performance. Our results provide compelling evidence suggesting targeted interventions to reduce abnormally high limbic coactivation may be an intriguing clinical application to ameliorate cognitive dysfunction associated with Alzheimer’s disease, which should be an active focus of future research.

## METHODS

### Data Source

Data used in the preparation of this article were obtained from the Alzheimer’s Disease Neuroimaging Initiative (ADNI) database (adni.loni.usc.edu). The ADNI was launched in 2003 as a public-private partnership, led by Principal Investigator Michael W. Weiner, MD. The primary goal of ADNI has been to test whether serial magnetic resonance imaging (MRI), positron emission tomography (PET), other biological markers, and clinical and neuropsychological assessment can be combined to measure the progression of mild cognitive impairment (MCI) and early Alzheimer’s disease (AD). Ethical approval was obtained by the ADNI investigators at each participating ADNI site in accordance with relevant guidelines and regulations, and all participants provided written informed consent.

### Structural and Functional MRI Acquisition and Processing

Structural and functional MRI data was downloaded in January 2023. As of that date, 405 advanced protocol resting state functional MRI scans (“Axial MB rsfMRI”) were identified. 34 scans were excluded after downloading due to inconsistent acquisition parameters (n=28; ADNI sites 037 and 177) or incomplete acquisitions (n=6), resulting in a total of 371 scans meeting criteria for preprocessing and quality control. Included scans had the following acquisition parameters: 3T Siemens Prisma or Prisma Fit; TR/TE 607/32 ms; 2.5mm^3^ resolution; flip angle = 50 degrees, 976 volumes; 704−704 matrix. The associated T1-weighted magnetization prepared rapid acquisition gradient echo (MPRAGE) scan collected at the same session as rsfMRI was also downloaded. T1 scans had the following acquisition parameters: echo time (TE) = 2.98 ms, repetition time (TR) = 2300 ms, inversion time = 900ms, flip angle = 10◦, field of view (FOV) = 208×240×256 mm3, acquired resolution = 1×1×1 mm. Additional information on MRI protocols can be found at: http://adni.loni.usc.edu/methods/documents/mri-protocols/.

Preprocessing of functional and structural images was performed using the CONN toolbox^68^ (version 21a) implemented in MATLAB version 2019b (MathWorks). Structural MRI was first segmented into gray matter (GM), white matter (WM), and cerebrospinal fluid (CSF) compartments. Structural MRI was then normalized to 1mm MNI space, and the resulting transformations were applied to GM/WM/CSF segments. Resting state functional data was realigned and coregistered to the T1 structural image, and the structural normalization parameters were applied to obtain 2mm MNI space functional images. No spatial smoothing was applied to preserve the high spatial resolution of the functional data.

All structural and functional images were intensively quality checked. The ART REPAIR toolbox, implemented in CONN, was used to detect outlier frames in the functional scans using a <0.5mm motion threshold and z-score >3. We excluded subjects who had outlier volumes greater than 2 standard deviations above the median value of the sample (threshold of 28.86% outlier volumes; median = 2.46%, SD = 13.2%). This resulted in the exclusion of 28 scans (7.5% of sample). See **Extended Data Figure 9** and **Extended Data Table 4** for the distribution of outlier volumes and framewise displacement^69^ in included and excluded subjects. Of the remaining scans, 4 were excluded for poor scan quality (e.g. incomplete field of view, artifacts) and 5 were excluded due to poor coregistration and warp quality. This resulted in a final analysis sample of 334 rsfMRI scans corresponding to 201 unique participants.

Mean BOLD time series were extracted from 218 regions of interest (ROIs) from the Brainnetome Atlas^70^. We included all cortical ROIs, as well as subcortical structures such as hippocampus and amygdala that are critical to memory and Alzheimer’s disease. Template space Brainnetome ROIs were masked with subject-specific gray matter masks prior to times series extraction to ensure signal was contained within gray matter.

Denoising was then performed on the 334 functional scans using CONN. First, despiking was applied to minimize effects of large spikes due to motion or other confounding factors. Despiking, rather than other methods such as scrubbing^69^, was applied because dynamic functional connectivity analyses require continuous samples for temporal sequencing analyses such as transition probabilities. The effects of six realignment parameters and their first-order derivatives (translations and rotations), anatomic CompCor^71^ (first five components of time series signal from white matter and CSF), and the potential “ramp up” effects at the beginning of the scan were regressed from the time series, and linear detrending was then applied to the residual time series. Finally, a bandpass filter (0.008-0.1 Hz) was applied after regression using a Fast Fourier Transformation (FFT) based procedure.

### Brain State Cluster Identification

The identification of brain states and quantification of brain state features and transition probabilities was performed using code based upon Cornblath et al. 2020^2^ (available at https://github.com/ejcorn/brain_states).

First, the denoised mean BOLD signal at each temporal volume (TR) for each scan was extracted for each of the 218 ROIs and entered into a data matrix, resulting in a 325,984 (334 scans x 976 temporal volumes) by 218 (ROIs) matrix.

To facilitate initialization of centroids, we created an exemplar matrix with about 5% of volumes identified to have the highest variance across ROIs for each subject. For each scan, volumes reflecting local maxima in variance (“peaks”) were identified from the full sample of volumes. We then removed successive peaks (i.e. two volumes in a row) to prevent redundancy and increased autocorrelation, and peaks that corresponded to a volume with a framewise displacement^69^ value over 0.25mm. This process resulted in 4.63% of observations within the initial data matrix being selected for the exemplar matrix, consistent in value with previous work^72^. This exemplar matrix was entered into k-means clustering algorithm (random centroid initialization, 20 replications, 500 iterations each for convergence), and the resulting cluster medians were entered as the initial centroids for full k-means clustering.

Next, k-means clustering was applied to the full data matrix using the cluster medians from the exemplar matrix for centroid initialization (*k*=2 to *k*=10, correlation distance metric, 20 replications, 500 iterations each for convergence). For each level of *k*, the variance explained (between cluster variance / [between within cluster variance + within cluster variance]) and silhouette scores^73,74^ were calculated. For additional validation, we performed sub-sample validation on the centroids for each level of *k*. We randomly sub-sampled 80% of the full dataset 500 times and submitted these subsampled matrices to k-means clustering to quantify the percentage of volumes that were assigned to the same brain state as in the original solution^75^ to assess reliability.

The optimal number of clusters was picked by considering a number of factors: variance explained, within cluster consistency (silhouette score), consensus clustering (reliability across random and unique data sub-samples), and ensuring no absent states in any of the scans. Elbow criterion were used to determine the highest *k* level before any additional clusters accounted for less than 1% variance explained gain^2,76^ (<1% gain from cluster *k* to cluster *k+1)*. Silhouette scores were used to quantify the degree of similarity between datapoints and their assigned cluster compared to neighboring clusters, with values approaching 1 indicating high similarity to the assigned cluster, and approaching −1 indicating high similarity to the neighboring clusters. We also considered consensus clustering score to ensure reliability of clusters at each level of *k*. Finally, to enable comparison of brain state dynamics across subjects, we ensured the selected *k* value allowed each subject to experience each brain state at least once (i.e., no absent states across subjects). Based on these criteria, *k*=6 was selected for subsequent analyses (71.5% variance explained). The transition from *k=6* to *k=7* resulted in an <1% gain in variance explained (0.07%)^2^. Additionally, *k=6* demonstrated a high silhouette score (mean = 0.064, SD = 0.052) compared to additional levels of *k*, a high average subsample validation score (99.45%), and no absent states across scans (see **Extended Data Figure 1**).

### Brain State Cluster Validation

Using *k=6* clusters, we next performed rigorous validation of the real clusters compared to null data models. Surrogate matrices for null hypothesis testing were generated for the following control models (**Extended Data Figure 2a**): (1) Independent Phase Randomization (IPR) model, which preserves the autocorrelation within each region but destroys covariance between regions^2,77^; (2) Iterative Amplitude Adjusted Fourier Transform (iAAFT) model, which preserves both the linear structure and the amplitude distribution autocorrelation function, resulting in equivalent second-order properties to the original data^78^, and (3) Random Gaussian model, which generates values from a normal distribution with the same mean and standard deviation as the real data.

For each null data model, k-means clustering was performed using *k*=6 and equivalent parameters as the real data (null data exemplar matrix centroid initialization, 20 replications, 500 iterations each for convergence). Silhouette scores were calculated for each null data model (**Extended Data Figure 2b**). Silhouette scores of the real data were significantly higher compared to IPR (IPR silhouette score = 0.031 ± 0.030; t(651,966) = 310.45, p = 0, CI = [0.0323-0.0327]), iAAFT (iAAFT silhouette score = 0. 0.061 ± 0.05; t(651,966) = 20.42, p = 1.22e^−92^, CI = [0.0023-0.0028]), and Random Gaussian models (Random Gaussian silhouette score = 0.005 ± 0.0004; t(651,966) = 650.28, p = 0, CI = [0.0588-0.0592]). These comparisons are visualized in **Extended Data Figure 2c**. These results support the validity of the real clustering solution.

We also tested the effect of removing high motion frames (framewise displacement >0.25 mm) in generating the real centroids for *k=6*, and found that the spatial correlation between the original centroids and the motion scrubbed centroids was *r* >0.99, indicating that the presence of high motion frames did not influence the centroid selection. We next also tested the effect of removing the initial 10 frames of the scan (∼1%) on centroid selection to correct for any potential “ramping up” effects of the scanner. This control similarly resulted in an *r* >0.99 spatial correlation between original and initial-frame dropped centroids. Finally, we conducted split half validation of our sample. Data was split into equal training and test groups and k-means clustering was run on each subsample. This process was iterated 500 times. This control resulted in *r* >0.99 median spatial correlation between original and split-half generated centroids (see **Extended Data Figure 3a**). Split half validation was also applied to test consistency of transition probability (median *r* >0.98) and persistence probability (median *r* = 0.72; see **Extended Data Figure 3b-c**).

### Brain State Persistence and Transition Probability Analyses

As part of the k-means clustering solution, each volume (TR) for each scan was assigned into the most similar brain state cluster. Using this information, we calculated three state persistence metrics (**Figure 1c; Extended Data Figure 4**): (1) fractional occupancy, or the percentage of volumes that were assigned to each state across the scan; (2) appearance rate, or the average rate at which the state appeared per minute; and (3) dwell time, or the average continuous time spent in a state when it appeared (in seconds). This analysis resulted in three metrics per state, for a total of 18 persistence features. Group-level distributions of values as well as persistence feature correlations across states are depicted in **Extended Data Figure 4**.

To determine whether there was a meaningful pattern of brain state persistence features across scans, we performed another level of clustering on the 18 persistence features. Each persistence feature was first normalized to enable comparisons across the different metrics (fractional occupancy, dwell time, appearance rate). Next all 18 persistence features were entered into a k-means algorithm (random centroid initialization, *k*=2 to *k*=8, Euclidean distance metric, 20 replications, 500 iterations each for convergence). Consistent with the initial brain state clustering, we derived variance explained and performed consensus clustering on the resulting clusters (**Extended Data Figure 5a**). Based upon these criteria, *k=4* was chosen as the ideal number of profiles. To validate our profile analyses, we reran clustering using a Gaussian Mixture Model approach with *k=4*, which resulted in highly similar patterns of profiles to the original k-means approach **Extended Data Figure 5b-c**). Further, to compare our results to a null model, persistence data was shuffled, and k-means clustering was performed on the shuffled data using the same parameters as the original k-means approach. This control resulted in profiles that were not distinguishable from each other (**Extended Data Figure 5d**), confirming that the k-means clustering performed on the real data was capturing real variance in the data.

Transition probabilities between brain states were calculated by comparing the probability of transition from a state at time *t* to a state at time *t+1*^2^. First, we calculated the transition probabilities using the full temporal sequence of states represented at each timepoint, which demonstrated that states are likely to persist rather than change to a distinct state. Second, we removed repeating states from the temporal sequence to calculate the probability of transitioning from one state to another distinct state to control for effects of state persistence^2^. Because this second approach better characterized trajectories of state transitions, this was used in our primary analyses to relate state transitions to other outcome measures. Normalized mutual information, reflecting how much information the current state holds in predicting the subsequent state, was calculated by comparing NMI between the original brain state time series and a brain state time series lagged by one element^2^. This was performed on the brain state time series with repeated states removed to control for effects of state persistence and focus on trajectories between distinct states.

### Control LIM State Analyses

Due to the spatial pattern of the LIM brain states overlapping with regions known to be vulnerable to signal drop out in fMRI, we investigated this possibility to ensure it was not influencing our results. For the LIM+ brain state, characterized by higher amplitude coactivation in anterior and medial temporal regions, we identified the top contributing Brainnetome ROIs with positive signal (23 ROIs or about the top 10%). Using these ROIs, we extracted the mean signal of each participant’s preprocessed functional image, masking the LIM ROIs with subject specific gray matter masks that were used to obtain the regional signal used in the brain states analysis.

LIM+ state fractional occupancy was correlated with the mean signal across the strongest LIM state contributor ROIs (r = −0.22, p<0.001), with higher LIM signal associated with less time spent in the LIM+ state. However, participants with the lowest signal did not overwhelmingly have high LIM+ fractional occupancy values. Within those with LIM+ mean signal below the median value, there was no correlation with LIM+ FO (n=166; r = −0.004, p = 0.96). This discrepancy does not provide convincing evidence that LIM+ FO is alone driven by signal drop out within these regions.

Further, to comprehensively reject the possibility that LIM signal drop out is driving our primary results, we tested whether our major outcome measures (age, clinical status, memory, etc.) were associated with LIM signal, and if controlling for LIM signal affected correlations between variables of interest and LIM+ FO. First, LIM signal was not significantly associated with major outcome measures such as age (r = 0.02, p = 0.66), diagnosis (CN vs. MCI/dementia, t(332) = 0.48, p = 0.64), CDR-SB status (0 vs. >0; t(326) = −0.18, p = 0.86), or memory performance (r = −0.02, p = 0.74). Second, LIM+ FO was still significantly associated with these factors while controlling for LIM signal: age (r = 0.33, p<0.001), diagnosis (F(1) = 6.94, p = 0.009), CDR-SB (F(1) = 10.7, p = 0.001), and memory (r = −0.24, p<0.001). Taken together, this is sufficient evidence that our brain state results are not driven by signal drop out.

### Static Resting State Network Analysis

To better interpret brain state persistence features in the context of traditional measures of static functional connectivity (FC), we calculated within-network FC, between-network FC, and system segregation^33,79^ of each of the canonical Yeo resting state networks^41^. This was performed across ROIs previously identified as members of these networks (i.e. visual network, somatomotor network, dorsal attention network, ventral attention network, limbic network, frontoparietal network, default mode network), and was independent from regions identified in our brain state clustering solution. ROI-to-ROI static functional connectivity was calculated using the CONN toolbox and ROIs from the Brainnetome Atlas^70^, and the Fisher’s z-transformed correlation coefficient was extracted between each ROI pair. First, for each network, within-network FC was calculated by taking the mean correlation value across ROIs contained within that resting state network. Second, for each network, between-network FC was calculated by taking the mean correlation value between each ROI contained in that network with all ROIs not contained within that network. Finally, for each network, system segregation was calculated as the difference between the networks within-network FC and between-network FC, divided by within-network FC.

### Pathology and Neurodegeneration Data

To assess relationships between Alzheimer’s pathology and brain state characteristics, we obtained positron emission tomography (PET) data from ADNI3, using 18F-Flortaucipir (FTP) for tau pathology and either 18F-Florbetapir (FBP) or 18F-Florbetaben (FBB) for amyloid-beta pathology (depending on availability and time match to rsfMRI). All PET data was preprocessed by the ADNI PET Core at UC Berkeley. Full acquisition and processing details are available on ADNI website (https://adni.loni.usc.edu/wp-content/uploads/2012/10/ADNI3_PET-Tech-Manual_V2.0_20161206.pdf).

In brief, FTP data was analyzed between 80-100 min post-injection across four 5-minute frames, partial volume corrected using a modified Rousset approach, and normalized using an inferior cerebellum gray reference region^80–82^. The mean standardized uptake value ratio (SUVR) values of regions corresponding to the FreeSurfer atlas (version 7.1.1 processing) were used for analyses. For the sparse canonical correlation analysis, all bilateral cortical regions contained within Braak I-IV regions^83^ (excluding the hippocampus due to off-target binding effects, and thalamus) were used. For a priori analyses, we focused on the mean SUVR of the entorhinal cortex and temporal meta-ROI^42^ (composite of the entorhinal, parahippocampal cortex, amygdala, fusiform, medial temporal, inferior temporal).

FBP data was analyzed between 50-70 min post-injection, while FBB data were analyzed 90-110 min post-injection. both across four 5-minute frames. Both FBP and FBB were normalized using a whole cerebellum reference region. Global amyloid-beta was calculated using a cortical summary region consisting of Freesurfer-defined (version 7.1.1 processing) frontal, anterior/posterior cingulate, lateral parietal, and lateral temporal regions^43,44^. To enable combination of the two tracers, regional and global FBP and FBB values were converted to the centiloid scale, and a threshold of 18 centiloids was used on the cortical summary region to determine amyloid-beta positivity^84^. For the sparse canonical correlation analysis, all bilateral cortical regions contained within Braak I-IV regions were included to be consistent with the tau analyses, with the addition of the hippocampus, with values represented in centiloids. For *a priori* analyses, we focused on amyloid-beta positive versus negative status.

For analyses assessing neurodegeneration, we used mean bilateral volume values from the FreeSurfer Cross-sectional version 6.0 processing pipeline in ADNI. Regional values were normalized by intracranial volume prior to analyses. For the sparse canonical correlation analysis, all cortical regions contained within Braak I-IV regions were used to be consistent with the tau analyses, with the addition of the hippocampus. For a priori analyses, we focused on the hippocampal volume, as this measure is known to be a sensitive marker of AD-related neurodegeneration.

Each tau-PET and Aβ-PET scan were time-matched to the nearest rsfMRI scan. Each PET measurement was only matched with one rsfMRI session to ensure no duplicate values were included in the analyses. Further, we excluded PET scans collected over 18 months from the rsfMRI visit. This resulted in 206 tau-PET/rsfMRI matches (mean 60 ± 98.3 days from rsfMRI) and 228 Aβ-PET/rsfMRI matches (mean 56 ± 94.4 days from rsfMRI). Direct correlations between state persistence features and PET measures controlled for rsfMRI-PET time interval. Freesurfer volumes were restricted to the same session as rsfMRI, resulting in 285 volume/rsfMRI matches used in analyses for the respective domains.

### Clinical and Cognitive Data

Clinical and cognitive data was obtained from ADNI3. Diagnosis was based upon ADNI’s criteria of cognitively normal (cognitively unimpaired, CU), mild cognitive impairment (MCI), or dementia. Due to the relatively small proportion of dementia diagnoses in our sample (n=31, 9.3%), we combined MCI and dementia into an cognitively impaired (CI) group. To characterize clinical impairment, we used the Sum of Boxes score from Clinical Dementia Rating (CDR-SB), which is a structured interview assessing cognitive, behavioral, and functional impairment across six domains: memory, orientation, judgment and problem solving, community affairs, home and hobbies, and personal care^85^. We dichotomized our sample into no clinical impairment (CDR-SB=0) versus any observable clinical impairment (CDR-SB>0). For another sensitive measure of clinical impairment that may better reflect the preclinical stage of Alzheimer’s disease, we used the Montreal Cognitive Assessment^86^ (MoCA).

To assess cognitive performance, we used composite scores based upon neuropsychological measures that were created for domains of episodic memory (“ADNI_MEM”) and executive function (“ADNI_EF”). Specific neuropsychological measures and methods for the calculation of the composite scores are detailed at: https://ida.loni.usc.edu/download/files/study/c042fcbc-61b5-402c-b20c-23892e3c8cd0/file/adni/ADNI_Methods_UWNPSYCHSUM_March_2020.pdf. To further probe episodic memory, we also used the immediate recall score from the Rey Auditory Verbal Learning Test^87^ (RAVLT), which is a word-list learning task that probes verbal memory that is commonly used across laboratories. Because this measure of memory is more specific for recall and episodic processes than some of the measures included in the ADNI memory composite, we replicated all associations with RAVLT immediate recall, which showed very similar, if not more robust, associations to the ADNI memory composite (e.g. profile comparison F(3) = 4.61, p = 0.004; see **Extended Data Figure 10** for all data).

### Statistical Analyses

Statistical analyses were conducted using MATLAB version 2019b, jamovi version 1.6.23.0, and R version 4.0.4. Correlations between factors were performed using Pearson’s r, and differences between groups were performed using independent samples t-tests for continuous variables and Chi-square analysis for discrete variables. Differences in outcomes between brain state profiles were tested using ANOVA models, testing for a main effect of profile (p<0.05). Planned follow-up pairwise comparisons were performed for significant ANOVA models, using p<0.05 (two-tailed) for post-hoc significance. Multiple comparison corrections were applied to the brain state transition probabilities (30 comparisons per matrix) and for comparisons of static FC (42 corrections per matrix) with the Benjamini & Hochberg FDR-correction^88^, using the p.adjust function from the “stats” package in R. Sparse canonical correlation analyses were performed using the “Penalized Multivariate Analysis (PMA)” R package^89^, using a 50% sparsity threshold and restricting PET values as positive loadings, and volume values as negative loadings to increase interpretability^90^. The first dimension was selected for interpretation. Mediation models, performed in jamovi, were considered significant when confidence intervals did not cross zero.

Data visualization was performed with Matlab and R. Brain state renderings were performed using Surf Ice (https://www.nitrc.org/projects/surfice/). Sparse canonical correlations were visualized with the “ggseg” package in R. General plots were created using “ggplot” in R. State transition pathways were created with the “qgraph” package in R. Figure panels 1c and 2d were created with Biorender.com, while all other figures were assembled with Adobe Illustrator.

## DATA AVAILABILITY

The data that support the findings of this study are available from The Alzheimer’s Disease Neuroimaging Initiative (ADNI) database (adni.loni.usc.edu) upon registration and compliance with the data usage agreement.

## CODE AVAILABILITY

The code generated during the current study is available from the corresponding author on reasonable request.

## ACKNOWLEGMENTS

Data collection and sharing for this project was funded by the Alzheimer’s Disease Neuroimaging Initiative (ADNI) (National Institutes of Health Grant U01 AG024904) and DOD ADNI (Department of Defense award number W81XWH-12-2-0012). ADNI is funded by the National Institute on Aging, the National Institute of Biomedical Imaging and Bioengineering, and through generous contributions from the following: AbbVie, Alzheimer’s Association; Alzheimer’s Drug Discovery Foundation; Araclon Biotech; BioClinica, Inc.; Biogen; Bristol-Myers Squibb Company; CereSpir, Inc.; Cogstate; Eisai Inc.; Elan Pharmaceuticals, Inc.; Eli Lilly and Company; EuroImmun; F. Hoffmann-La Roche Ltd and its affiliated company Genentech, Inc.; Fujirebio; GE Healthcare; IXICO Ltd.; Janssen Alzheimer Immunotherapy Research & Development, LLC.; Johnson & Johnson Pharmaceutical Research & Development LLC.; Lumosity; Lundbeck; Merck & Co., Inc.; Meso Scale Diagnostics, LLC.; NeuroRx Research; Neurotrack Technologies; Novartis Pharmaceuticals Corporation; Pfizer Inc.; Piramal Imaging; Servier; Takeda Pharmaceutical Company; and Transition Therapeutics. The Canadian Institutes of Health Research is providing funds to support ADNI clinical sites in Canada. Private sector contributions are facilitated by the Foundation for the National Institutes of Health (www.fnih.org). The grantee organization is the Northern California Institute for Research and Education, and the study is coordinated by the Alzheimer’s Therapeutic Research Institute at the University of Southern California. ADNI data are disseminated by the Laboratory for Neuro Imaging at the University of Southern California.

The opinions and assertions herein do not necessarily reflect the official policy or position of the Uniformed Services University, the Department of Defense or the Henry M. Jackson Foundation for the Advancement of Military Medicine. No conflicts of interest are reported.

## AUTHOR CONTRIBUTIONS

Conceptualization, J.N.A. & M.A.Y; Methodology & Software, J.N.A., S.M.K., M.G.C.F., Y.E., L.A.S., P.E.R.; Formal Analysis, J.N.A.; Writing - Original Draft, J.N.A.; Writing - Reviewing and Editing, all authors.

## EXTENDED DATA

**EXTENDED DATA FIGURE 1.**
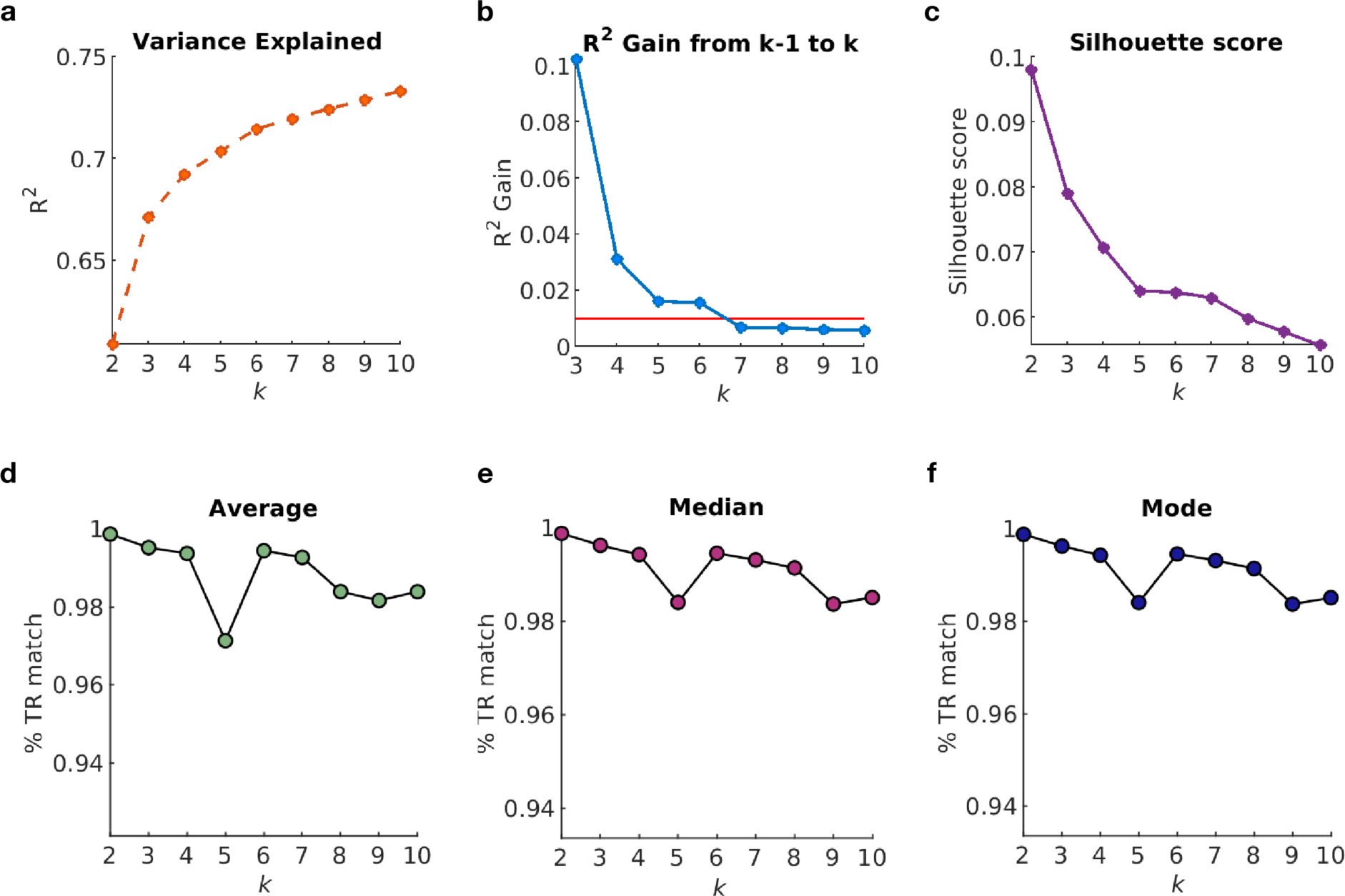
Criteria used for selection of *k* brain state clusters. (**a**) Elbow plot showing the total variance (R^2^) explained for each clustering solution. (**b**) Plot showing change in variance explained with progression to a higher k-value. Red line indicates 1% threshold. Moving from *k*=6 to *k*=7 results in <1% additional variance explained. (**c**) Silhouette score for each clustering solution, with higher scores indicating more similarity to the assigned cluster compared to neighboring clusters. (**d-f**) Consensus clustering was performed to determine subject validation of clustering solution across all levels of *k*. 80% of the full dataset was randomly sub-sampled 500 times and submitted to *k*-means clustering to quantify the percentage of frames (TRs) that were assigned the same brain state in the original solution and in the random sub-sample. The average (**d**), median (**e**), and mode (**f**) of the consensus clustering is shown for each level of *k*, with *k*=6 demonstrating a high % TR match.

**EXTENDED DATA FIGURE 2.**
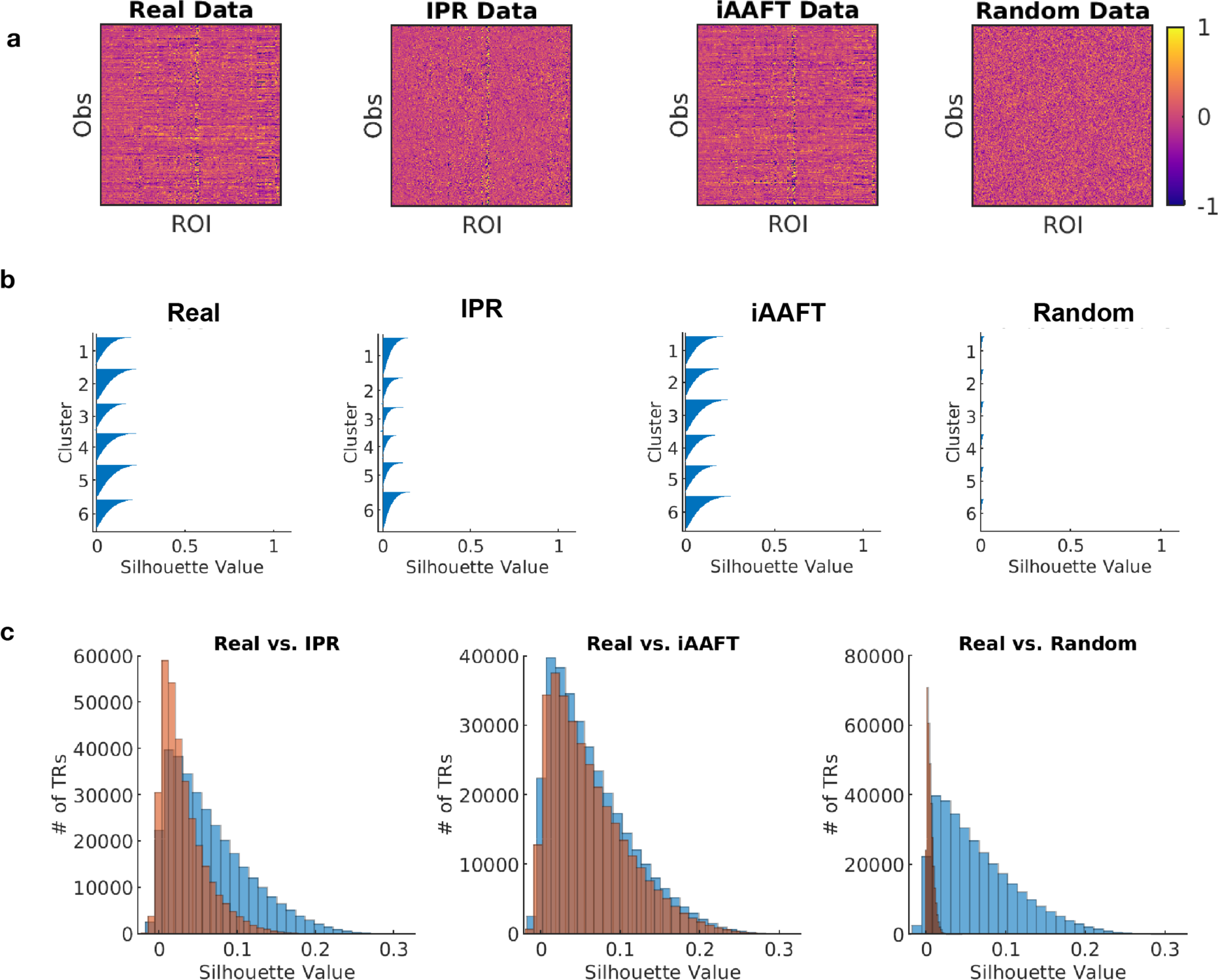
Comparison of real data and clusters to null surrogate datasets and clusters. (**a**) Surrogate datasets were created for null hypothesis testing with the following models: (1) Independent Phase Randomization (IPR) model, which preserves the autocorrelation within each region but destroys covariance between regions; (2) Iterative Amplitude Adjusted Fourier Transform (iAAFT) model, which preserves both the linear structure and the amplitude distribution autocorrelation function, resulting in equivalent second-order properties to the original data, and (3) Random Gaussian model, which generates values from a normal distribution with the same mean and standard deviation as the real data. (**b**) Silhouette values for k=6 clusters for the real data and each surrogate dataset. (**c**) Comparison of all silhouette values for real data (blue) compared to the IPR, iAAFT, and Random Gaussian clusters (orange). The real data had significantly higher (better) silhouette values compared to each null model.

**EXTENDED DATA FIGURE 3.**
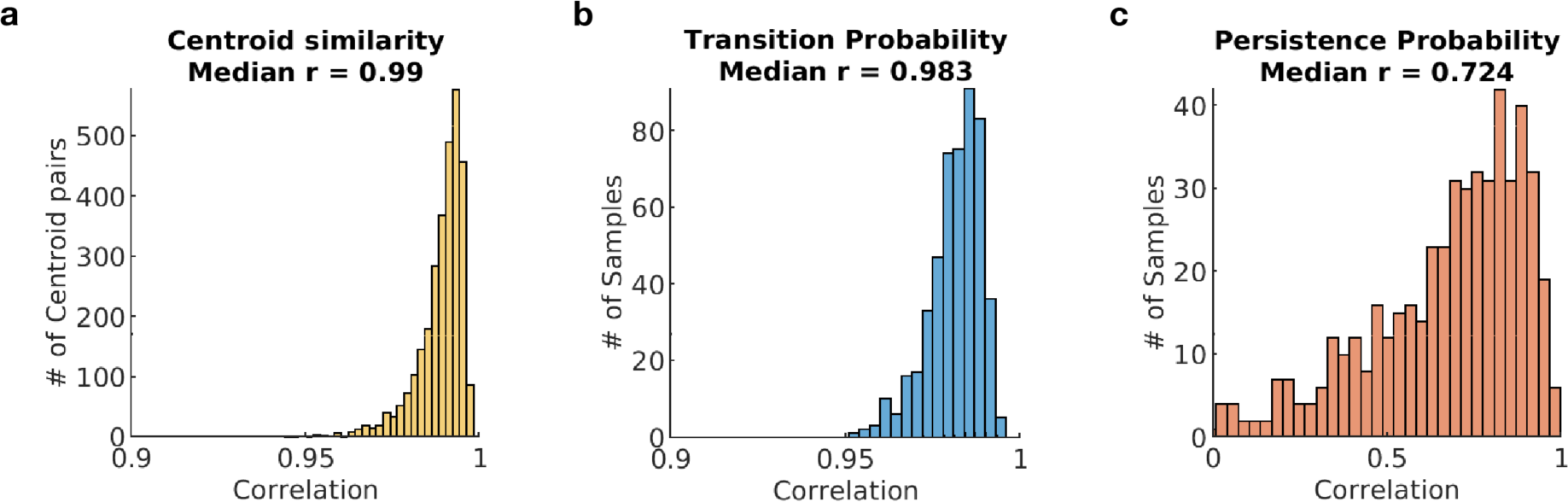
Split-halves validation of the chosen *k* (*k* = 6). The real data matrix of the study sample was randomly split in half (n=167 for training data, n=167 for test data) 500 unique times. K-means clustering was run separately for each half and correlation was used to quantify similarity between the centroid locations (**a**), transition probabilities (**b**), and persistence probabilities (**c**).

**EXTENDED DATA FIGURE 4.**
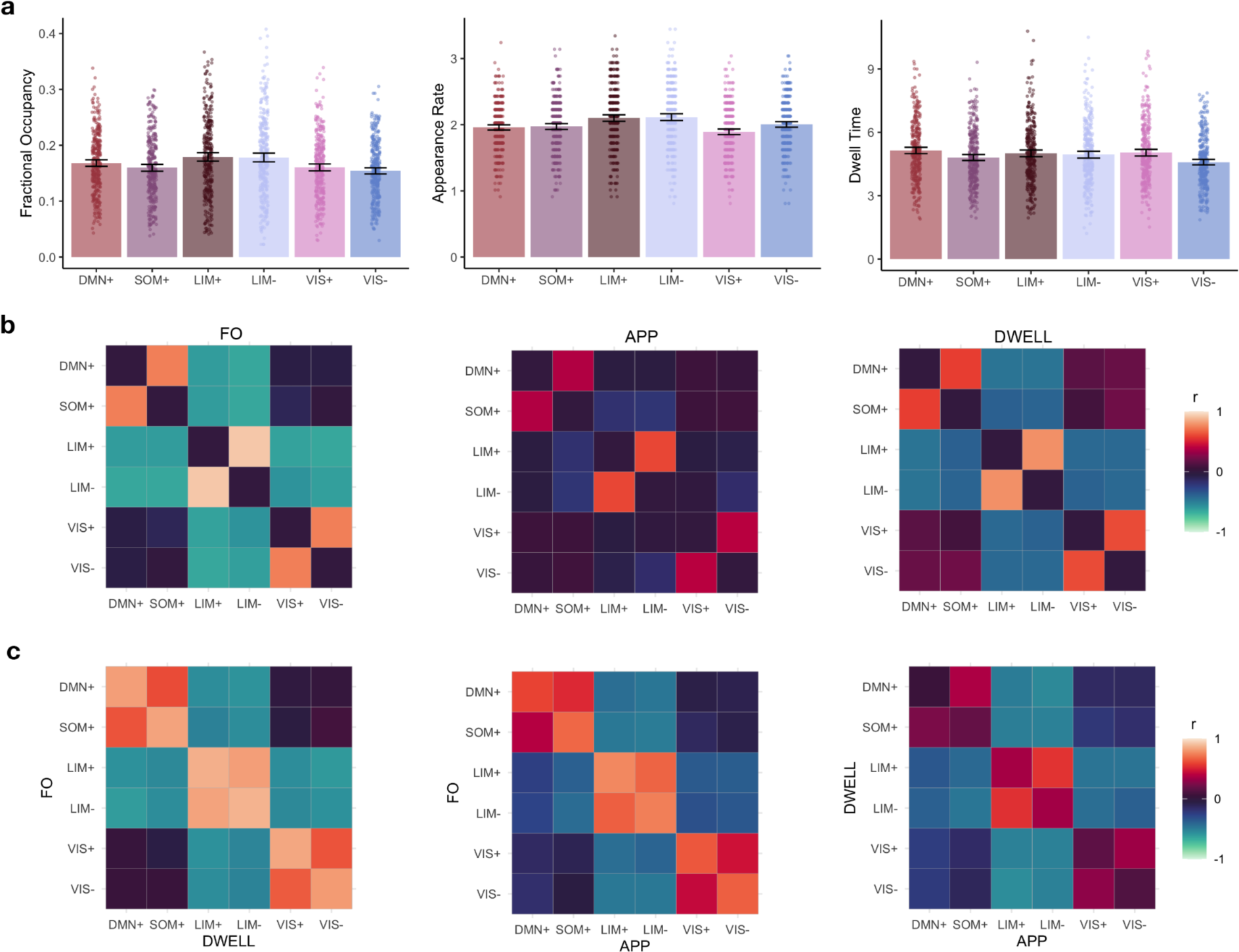
Group-level brain state persistence features. (a) Distributions of values for fractional occupancy (left), appearance rate (center), and dwell time (right). **(b)** Correlations between state features for fractional occupancy (left), appearance rate (center), and dwell time (right). **(c)** Cross-correlations between fractional occupancy and appearance rate (left), fractional occupancy and dwell time (center), and appearance rate and dwell time (right). FO, fractional occupancy; APP, appearance rate; DWELL, dwell time; DMN+, default mode network high amplitude coactivation state; SOM+, somatomotor high amplitude coactivation state; LIM+, limbic high amplitude coactivation state; LIM-, limbic low amplitude coactivation state; VIS+, visual high amplitude coactivation state; VIS-, visual low amplitude coactivation state.

**EXTENDED DATA FIGURE 5.**
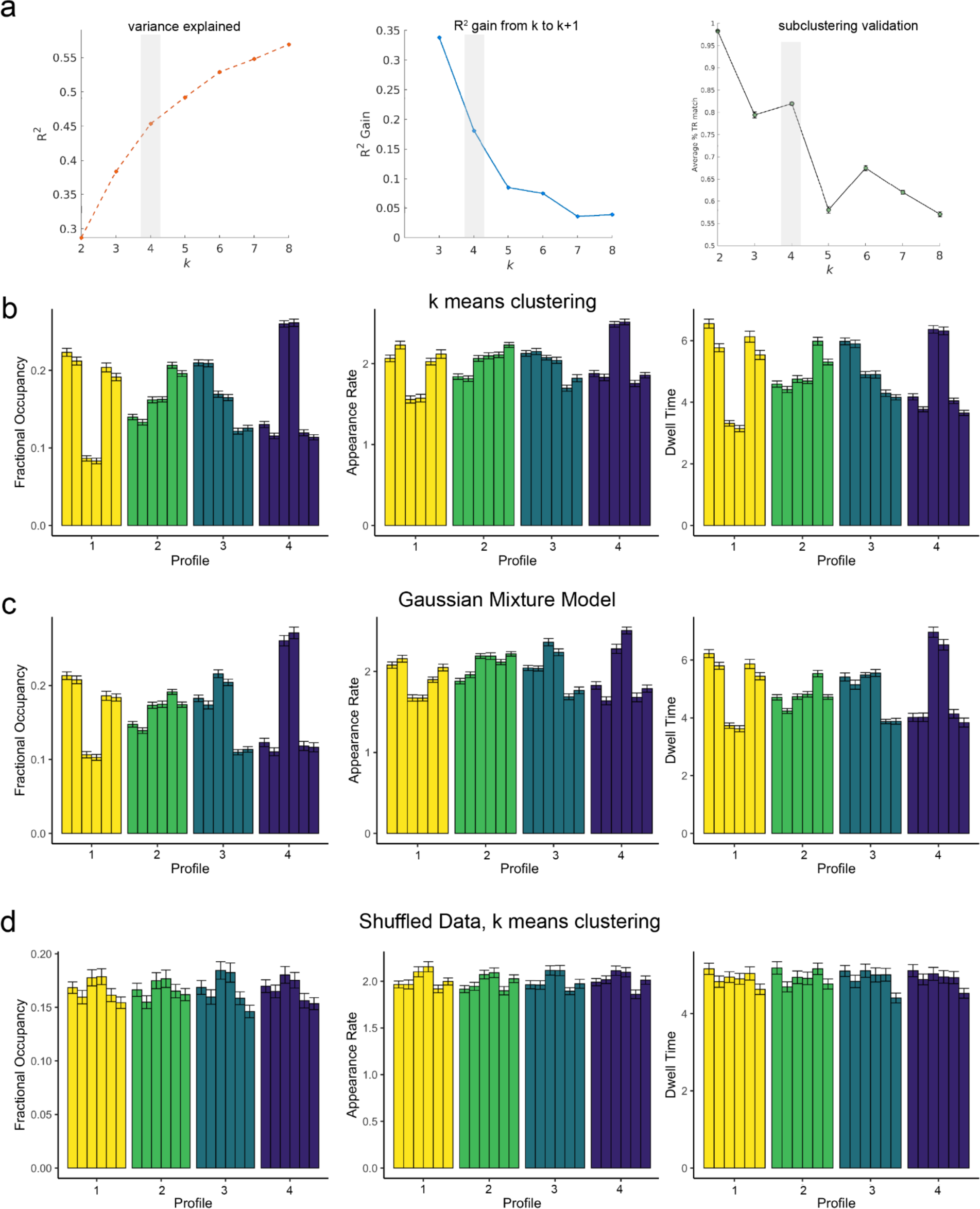
Selection and control models for brain state profiles. (a) Based upon converging results the inflection point of the elbow plot (left), variance (R2) gained from *k* to *k*+1 (center), and subclustering validation (right), *k*=4 was chosen as the optimal number of profiles (shaded region). **(b)** Results of k-means clustering (*k*=4), demonstrating distinct patterns of fractional occupancy (left), appearance rate (center), and dwell time (right) of the profiles. **(c)** Clustering solution of a Gaussian Mixture Model using four clusters revealed very similar patterns of profiles to the k-means clustering approach. **(d)** Shuffled data applied to k-means clustering (*k*=4) does not demonstrate dissociable patterns of persistence features between profiles.

**EXTENDED DATA FIGURE 6.**
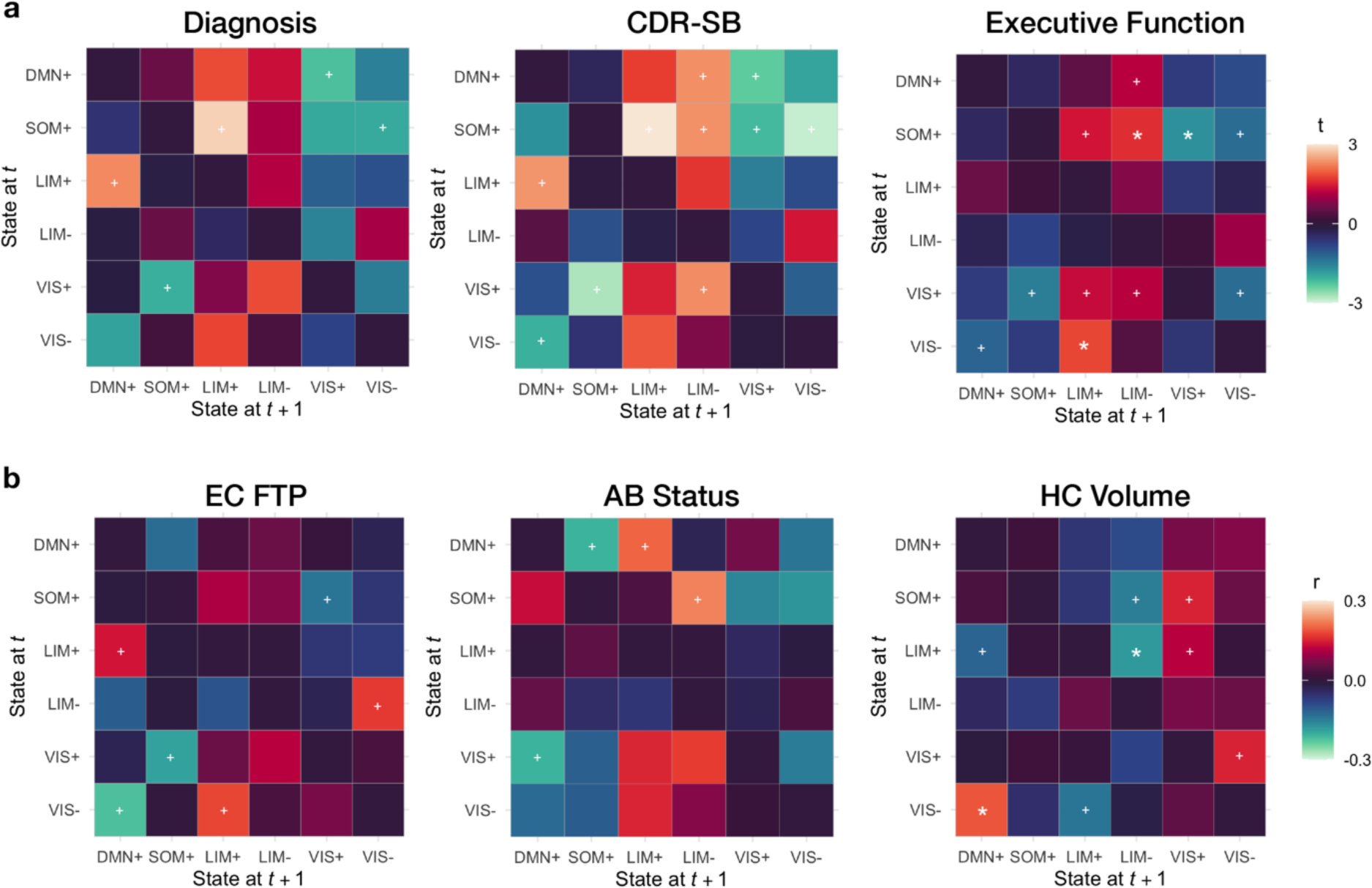
Additional associations with brain state transitions. Associations between state transition probabilities with (**a**) clinical and cognitive features and (**b**) Alzheimer’s neuropathology. Correlation coefficients are represented as positive (reds) or negative (blues) between each transition probability and outcome measure. Significance was corrected for multiple comparisons using FDR correction over 30 possible transitions (ignoring autocorrelation). White stars represent pFDR <0.05, and white crosses represent punc <0.05.

**EXTENDED DATA FIGURE 7.**
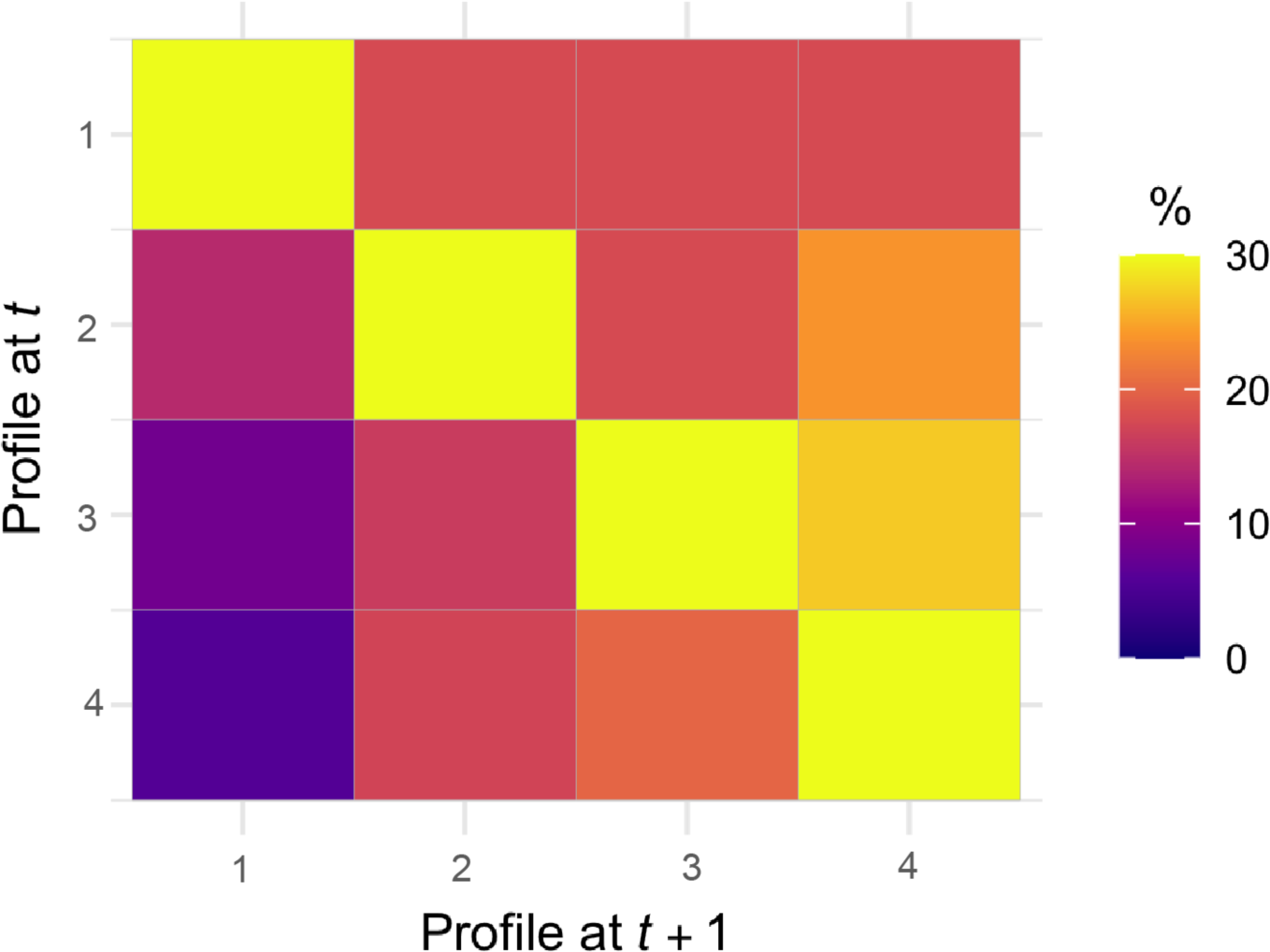
Stability and transitions between profiles over time. The probability of transitioning from a Profile at time point *t* (y-axis) to either the same or different Profile at time point *t + 1* (x-axis). Stability of profile transitions was most likely across all Profiles. Profile 1 (the healthiest profile) was equally likely to transition to any other profile, while Profile 4 (the unhealthiest profile) was very unlikely to transition to Profile 1 (the healthiest profile).

**EXTENDED DATA FIGURE 8.**
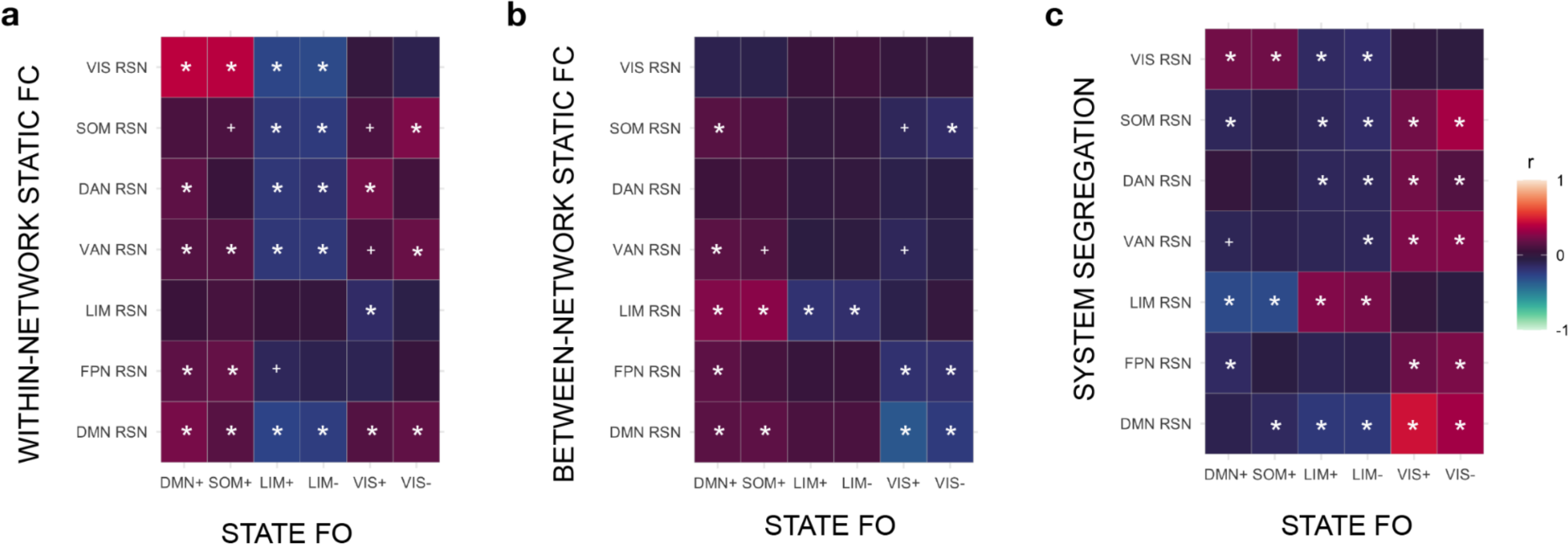
Associations between brain state fractional occupancy and static measures of functional connectivity. Static functional connectivity (FC) was calculated between ROIs included in the Brainnetome Atlas. For each canonical Yeo resting state network (RSN), we calculated within-network FC (**a**), between-network FC (**b**), and system segregation (**c**). Associations between the fractional occupancy of each brain state and the static FC of the various RSNs are depicted as correlation matrices. Static RSNs: VIS RSN, visual network; SOM, somatomotor network; DAN, dorsal attention network; VAN, ventral attention network; LIM, limbic network; FPN, frontoparietal network; DMN, default mode network; Brain states: DMN+, default mode network high amplitude brain state; SOM+, somatomotor high amplitude brain state; LIM+, limbic high amplitude brain state; LIM-, limbic low amplitude brain state; VIS+, visual high amplitude brain state; VIS-, visual low amplitude brain state. White asterisks indicate pFDR <0.05 (corrected for 42 comparisons), while white crosses indicate punc <0.05.

**EXTENDED DATA FIGURE 9.**
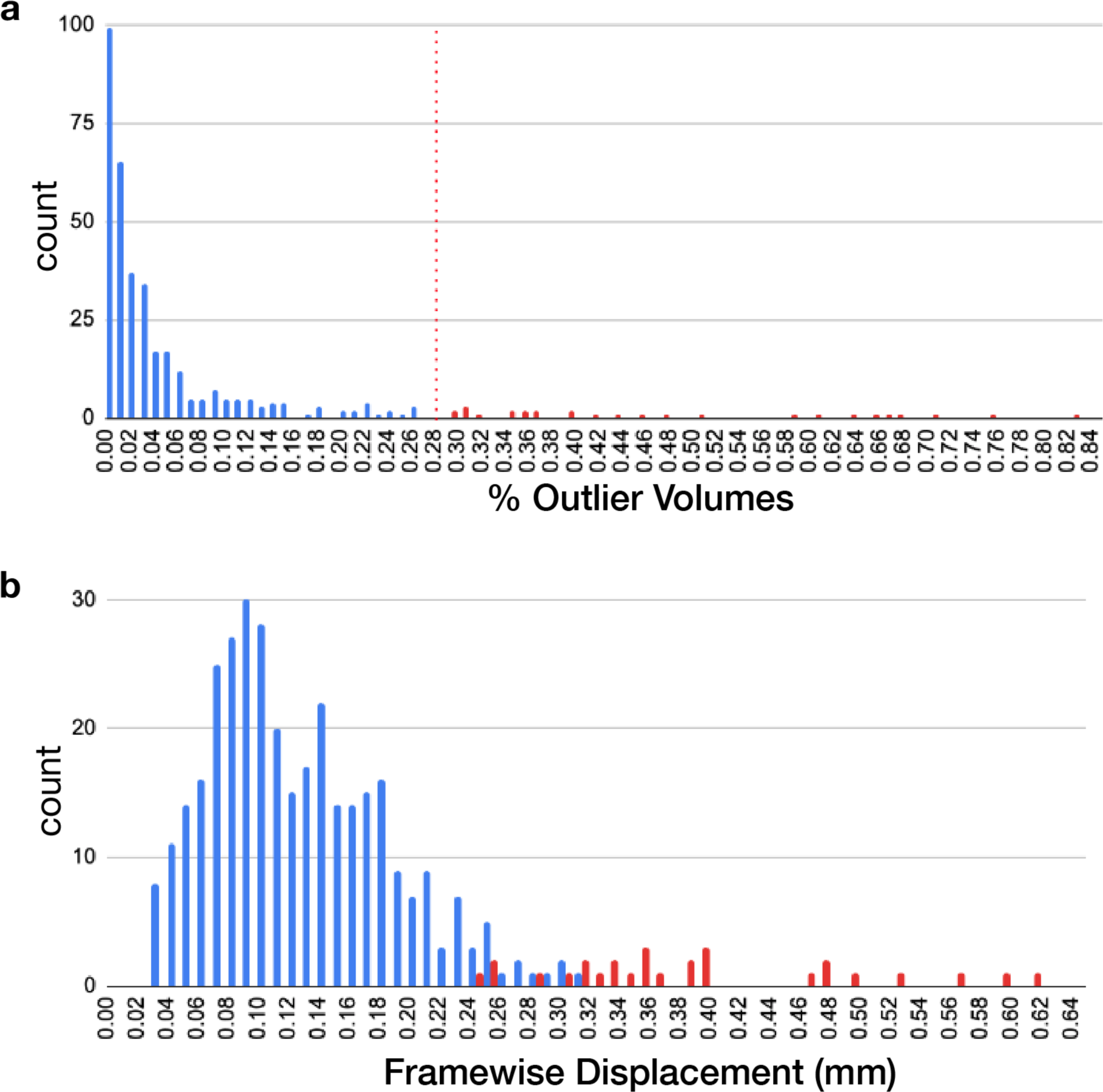
Distribution of motion and thresholds for exclusion. (**a**) Distribution of outlier volumes based upon a conservative threshold of <0.5mm framewise displacement and global z-score <3. The dotted red line indicates the exclusion threshold (<28.86%), calculated as two standard deviations above the median value. Red vales on the histogram represent scans excluded by falling over this threshold. (**b**) Distribution of framewise displacement (FWD) as calculated by Power et al., 2012. Red values on the histogram represented excluded scans based upon the outlier volume threshold applied in (**a**).

**EXTENDED DATA FIGURE 10.**
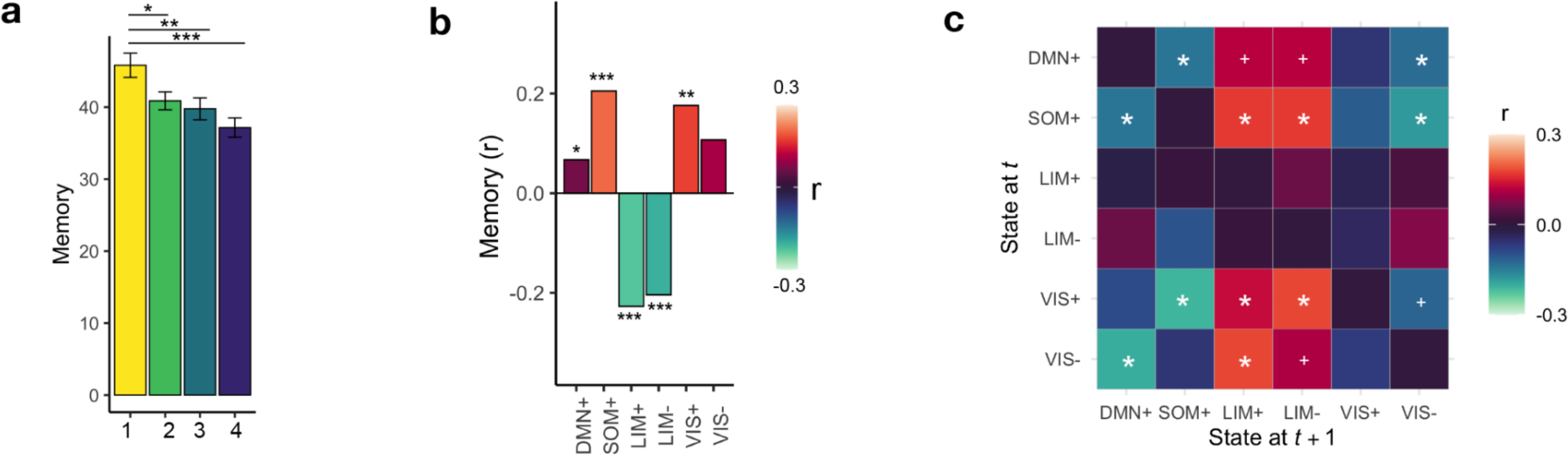
Replication of primary memory results using Rey Auditory Verbal Learning Test (RAVLT) immediate recall score. (**a**) Profile comparison indicating decreased RAVLT immediate score in all groups compared to Profile 1. (**b**) Direct correlations between fractional occupancy of each state and RAVLT immediate score. (**c**) Associations between state transition probabilities and RAVLT immediate score. White asterisks indicate pFDR <0.05 (for 30 comparisons), while white crosses indicate punc <0.05.

**EXTENDED DATA TABLE 1.** Demographics and key outcomes of the sample.

|  | Cognitively Unimpaired<br>(CU; n=188) | Cognitively Impaired*<br>(CI; n=146) | CU vs. CI |  |
| --- | --- | --- | --- | --- |
|  |  |  | t or X<sup>2</sup> | p |
| Age (years) | 72.8 (7.9) | 75.3 (8.0) | -2.85 | 0.005 |
| Sex (n, % female) | 115 (61.2%) | 55 (37.7%) | 18.2 | <0.001 |
| Education (years) | 16.8 | 15.7 | 3.94 | <0.001 |
| Race (n, % White) | 159 (84.6%) | 118 (80.8%) | 3.41 | 0.49 |
| Ethnicity (n, % Non-Hispanic/Latino) | 176 (93.6%) | 134 (91.8%) | 1.19 | 0.76 |
| CDR-SB | 0.05 | 2.14 | -13.58 | <0.001 |
| MMSE | 29.0 (1.11) | 26.7 (3.63) | 7.82 | <0.001 |
| MoCA | 26.0 (2.54) | 22.0 (4.30) | 9.03 | <0.001 |
| Memory Composite | 1.11 (0.77) | 0.16 (0.86) | 10.45 | <0.001 |
| Executive Function Composite | 1.12 (0.84) | 0.29 (0.95) | 8.17 | <0.001 |
| RAVLT Immediate Recall | 45.7 (11.8) | 33.5 (11.1) | 9.36 | <0.001 |
| EC FTP SUVR | 1.80 (0.34) | 2.31 (0.79) | -6.18 | <0.001 |
| MetaTemporal FTP SUVR | 1.59 (0.29) | 1.97 (0.72) | -5.11 | <0.001 |
| Aβ Status (% positive) | 39 (28.9%) | 56 (60.2%) | 22.20 | <0.001 |
| Global Aβ (centiloids) | 21.1 (31.2) | 46.3 (42.4) | -5.16 | <0.001 |
| HC Volume (cm3, normalized) | 2.51 (0.32) | 2.28 (0.34) | 5.98 | <0.001 |

\*CI group composed of patients with mild cognitive impairment (n=115) and dementia (n=31); Race, n=14 missing values; Ethnicity, n=17 missing values; Clinical Dementia Rating Sum of Boxes score (CDR-SB), n=6 missing values; Mini Mental State Exam (MMSE), n=15 missing values; Montreal Cognitive Assessment (MoCA), n=23 missing values; Memory Composite, n=15 missing values; Executive Function Composite, n=19 missing values; Rey Auditory Verbal Learning Test (RAVLT) immediate, n=17 missing values; 18F-Flortaucipir (FTP), n=128 missing values; EC, entorhinal cortex; MetaTemporal composite, composite of regions within the medial and lateral temporal lobe; SUVR, standardized uptake value ratio; Aβ , amyloid-beta as measured by 18F-florbetapir or 18F-florbetaben, n = 106 missing values; Aβ positivity defined as >18 centiloids; HC, hippocampus (normalized by intracranial volume), n=49 missing values.

**EXTENDED DATA TABLE 2.** Brain state profile clinical and cognitive comparisons.

|  | Profile 1 vs 2 |  | Profile 1 vs 3 |  | Profile 1 vs 4 |  | Profile 2 vs 3 |  | Profile 2 vs 4 |  | Profile 3 vs 4 |  |
| --- | --- | --- | --- | --- | --- | --- | --- | --- | --- | --- | --- | --- |
| | $t/X^2$ | $p$ | $t/X^2$ | $p$ | $t/X^2$ | $p$ | $t/X^2$ | $p$ | $t/X^2$ | $p$ | $t/X^2$ | $p$ |
| Age | -0.97 | 0.33 | **-3.46** | **<0.001** | **-4.81** | **<0.001** | **-2.84** | **0.005** | **-4.36** | **<0.001** | -1.24 | 0.22 |
| Dx | 0.01 | 0.93 | -3.47 | 0.06 | 2.98 | 0.09 | **-4.77** | **0.03** | **4.25** | **0.04** | 0.7 | 0.80 |
| CDR-SB | 0.99 | 0.32 | **-3.95** | **0.047** | **6.88** | **0.009** | -1.40 | 0.24 | 3.54 | 0.06 | 0.33 | 0.57 |
| MoCA | 1.46 | 0.15 | 1.31 | 0.19 | **3.27** | **0.001** | -0.10 | 0.92 | **1.99** | **0.047** | **2.00** | **0.046** |
| Memory | **1.97** | **0.049** | **2.61** | **0.009** | **3.63** | **<0.001** | 0.78 | 0.44 | 1.81 | 0.07 | 0.93 | 0.35 |
| EF | -0.02 | 0.98 | 0.71 | 0.48 | **2.78** | **0.006** | 0.80 | 0.43 | **3.11** | **0.002** | **2.15** | **0.03** |
| RAVLT | **2.33** | **0.02** | **2.75** | **0.006** | **4.15** | **<0.001** | 0.55 | 0.58 | **1.98** | **0.048** | 1.33 | 0.18 |

Dx, cognitively unimpaired versus cognitively impaired (mild cognitive impairment/dementia) diagnosis; Clinical Dementia Rating Sum of Boxes score (CDR-SB); Montreal Cognitive Assessment (MoCA); Memory, Memory Composite score; EF, Executive Function Composite score; Rey Auditory Verbal Learning Test (RAVLT) immediate score.

**EXTENDED DATA TABLE 3.** Results of linear and logistic regression analyses with LIM+ fractional occupancy as a predictor.

| **Model** |  |  |  |  |
| --- | --- | --- | --- | --- |
| **Memory** ($R^2 = 0.33$, $p < 0.001$) | **estimate** | **SE** | **t** | **p** |
| Intercept | 1.59 | 0.93 | 1.71 | 0.09 |
| Age | -0.006 | 0.008 | -0.79 | 0.43 |
| EC FTP | -0.68 | 0.10 | -7.12 | <0.001 |
| HC volume | 0.49 | 0.19 | 2.64 | 0.009 |
| LIM+ FO | -1.77 | 0.79 | -2.24 | 0.026 |
| **MoCA** ($R^2 = 0.38$, $p < 0.001$) | **estimate** | **SE** | **t** | **p** |
| Intercept | 36.02 | 4.18 | 8.61 | <0.001 |
| Age | -0.03 | 0.04 | -0.86 | 0.39 |
| EC FTP | -3.63 | 0.44 | -8.32 | <0.001 |
| HC volume | -0.40 | 0.84 | -0.47 | 0.64 |
| LIM+ FO | -8.79 | 3.52 | -2.49 | 0.01 |
| **CDR-SB** ($R^2 = 0.20$, $p < 0.001$) | **estimate** | **SE** | **t** | **p** |
| Intercept | 0.010 | 3.04 | 0.003 | 0.99 |
| Age | 0.03 | 0.03 | 0.96 | 0.34 |
| EC FTP | -1.51 | 0.38 | -3.94 | <0.001 |
| HC volume | 1.14 | 0.61 | 1.87 | 0.06 |
| LIM+ FO | -7.77 | 2.68 | -2.90 | 0.004 |
| **Diagnosis** ($R^2 = 0.17$, $p < 0.001$) | **estimate** | **SE** | **t** | **p** |
| Intercept | 0.16 | 2.96 | 0.06 | 0.96 |
| Age | -0.01 | 0.03 | -0.38 | 0.70 |
| EC FTP | 1.26 | 0.36 | 3.54 | <0.001 |
| HC volume | -1.25 | 0.60 | -2.08 | 0.04 |
| LIM+ FO | 4.56 | 2.53 | 1.80 | 0.07 |

Diagnosis, cognitively unimpaired versus cognitively impaired (mild cognitive impairment/dementia) diagnosis; Clinical Dementia Rating Sum of Boxes score (CDR-SB); Montreal Cognitive Assessment (MoCA); Memory, Memory Composite score; SE, standard error.

**EXTENDED DATA TABLE 4.**
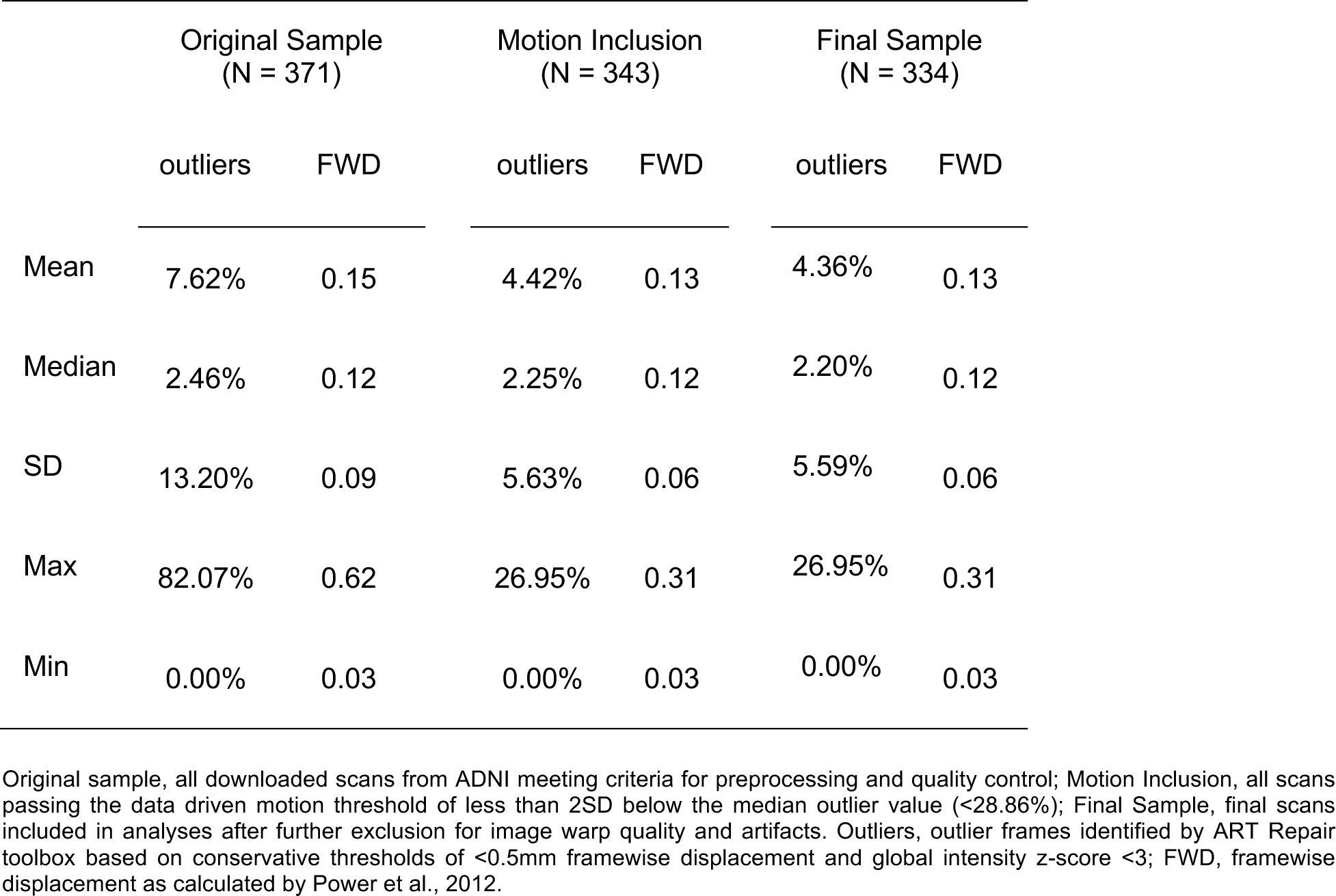
Motion outliers and framewise displacement.

